# RNA Binding of Syntaxin 5 Synergises Membrane Fusion to Accelerate miRNA Export, Activate Macrophage and Clear Pathogens

**DOI:** 10.1101/2023.09.08.556844

**Authors:** Sourav Hom Choudhury, Shreya Bhattacharjee, Kamalika Mukherjee, Suvendra N. Bhattacharyya

**Affiliations:** RNA Biology Research Laboratory, Molecular Genetics Division, CSIR-Indian Institute of Chemical Biology, Kolkata-700032, India; Academy of Scientific and Innovative Research (AcSIR), Ghaziabad-201002, India; Department of Pharmacology and Experimental Neuroscience, University of Nebraska Medical Center (UNMC), NE, USA

**Keywords:** miRNA export, STX5, SNARE protein, RNA binding protein, HuR

## Abstract

Intercellular miRNA exchange acts as a key mechanism to control gene expression post-transcriptionally in mammalian cells. Regulated export of repressive miRNAs allows the expression of inflammatory cytokines in activated macrophages. Intracellular trafficking of miRNAs from endoplasmic reticulum to endosomes is a rate determining step in miRNA export process and plays an important role in controlling cellular miRNA level and inflammatory processes in macrophages. We have identified the SNARE protein Syntaxin5 to show a synchronized expression pattern with miRNA activity loss in activated mammalian macrophage cells. Syntaxin 5 is both necessary and sufficient for macrophage activation and clearance of the intracellular pathogen *Leishmania donovani* from infected macrophages. Exploring the mechanism of how Syntaxin5 acts as an immunostimulant, we have identified the *de novo* RNA binding property of this SNARE protein that binds specific miRNAs and facilitates their accumulation in endosomes in a cooperative manner with human ELAV protein HuR to ensure export of miRNAs and allows the expression of miRNA-repressed cytokines. Conversely, in its dual role in miRNA export, this SNARE protein prevents lysosomal targeting of endosomes by enhancing the fusion of miRNA-loaded endosomes with plasma membrane to ensure accelerated release of extracellular vesicles and associated miRNAs.

**Graphical Abstract:** 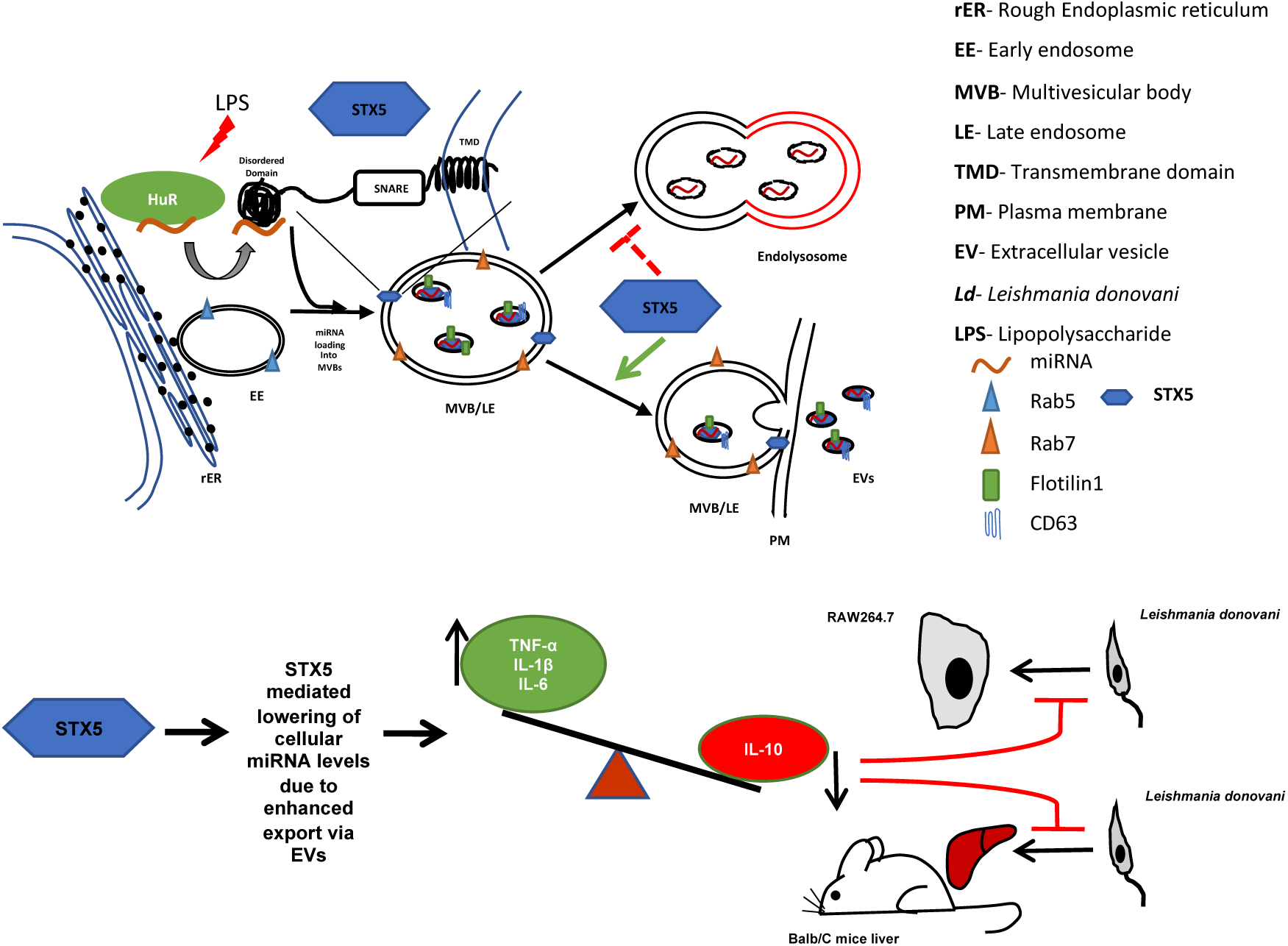

- *HuR transfers miRNAs to STX5 for export*
- *STX5 utilizes its N-terminal disordered domain to bind and target miRNAs to endosomes*.
- *In its dual role, STX5 uses the SNARE domain to export miRNA by promoting EV biogenesis and release*.
- *STX5-mediated miRNA export activates macrophage to clear internalized parasites*.

## Introduction

microRNAs (miRNAs) are 20-22 nucleotide long non-coding RNAs that orchestrate the complex network of post-transcriptional gene expression in metazoans^1,2^. The guide strand of miRNA is loaded into the RNA-induced silencing complex (RISC) to suppress gene expression by binding to the 3′ untranslated regions (3′-UTRs) of the cognate mRNAs, inducing mRNA degradation or translational repression upon target sequence recognition and base pairing by miRNAs^1-3^. miRNA-mediated repression and target mRNA degradation are spatio-temporally uncoupled events as the repression occurs on the surface of rough endoplasmic reticulum (rER) while the target RNA degradation takes place on late endosomes and multi vesicular bodies (MVBs)^4^. Overall, miRNA mediated gene regulation is a very intricate, organized, and compartmentalized process that occurs at different stages of translation repression happening at different subcellular compartments.

Other than the intracellular gene silencing, miRNAs also play a crucial role in intercellular communication as it has already been reported that hundreds of miRNAs are exported out of the animal cells via extracellular vesicles (EVs). MVBs fuse with the plasma membrane to release exosomes in the extracellular milieu as EVs and they contain several miRNAs, mRNAs, proteins and other cellular factors ^5,6^. Exosome bound extracellular miRNAs can be taken up by the neighbouring cells to regulate the gene expression while the miRNA exchange process gets affected in diseased tissue ^5,7,8^.

miRNAs also play a pivotal role in maintaining an optimal cytokine expression in macrophages in resting stage. Macrophage cells are usually able to clear out the invading pathogens by altering cytokine expression to upregulate the expression of pro-inflammatory cytokines^9,10^. Bacterial lipopolysaccharide (LPS) enhances the expression of pro-inflammatory cytokines by binding to the Toll-like receptor4 (TLR4) and affects the activity of different miRNAs in monocytes and macrophages^11^. Conversely, the protozoan parasite *Leishmania donovani*, the causative agent of visceral leishmaniasis, escapes this pro-inflammatory cytokine surge by inducing the expression of anti-inflammatory cytokines and inhibiting the lysosomal fusion of parasitophorous vacuoles inside the macrophages^9,12,13^.

EV biogenesis and the mechanism of miRNA packaging into those vesicles has been an important research area in the EV-research field. RNA binding proteins (RBPs) such as hnRNPA2B1, SYNCRIP, YBX-1, MVP and La protein specifically bind to their target miRNAs and play a crucial role in the sorting and packaging of miRNAs to exosomes^14-17^. Human ELAV protein HuR also plays a pivotal role in exporting out miRNAs via exosomes under cellular stress conditions^18^. HuR inhibits the cellular activity of specific miRNAs mainly by uncoupling them from their targets and by subsequently loading them inside into MVBs for their release as part of EVs ^19-21^.

On the other hand, EV biogenesis mainly occurs either by recruiting the ESCRT proteins or by synthesizing ceramide with the help of neutral sphingomyelinase 2 (nSMase2) ^22^. The exosomal release is a complex multistep process that is strictly regulated by ESCRT, Rab and other small GTPases and SNARE proteins^23,24^. The release of exosomes involves vesicle fusion, a process that is governed by several different SNARE proteins. There are reports that showed that even depletion of one specific SNARE proteins might lead to reduced exosome secretion in *C. elegans*^25^. However, how these SNARE proteins that modulate endosomal pathway affect the host-pathogen interaction by altering the cellular miRNA activity and levels to affect the target cytokine profile is yet to be explored in detail.

In this study, we have identified one t-SNARE protein, Syntaxin5 (STX5) that enhances the interaction between endoplasmic reticulum and early endosomes, which in turn facilitate the release of EVs. Interestingly, STX5 found to be essential for EV biogenesis and export pathways and thus to have its negative effect on cellular miRNA levels. Moreover, STX5 with a perturbed SNARE domain was unable to show its regulatory effect on cellular miRNA activity and level. Expression of STX5 induces a decrease in miRNA activity to allow expression of miRNA-repressed cytokines to restrict the *Leishmania donovani* infection in murine macrophage cells and in mice liver. We have also observed a *de novo* miRNA binding activity of STX5 both *in vivo* and *in vitro*. The N-terminal 23 amino acids long disordered region of STX5 found to essential for miRNA binding and HuR interaction. Interestingly, human ELAV protein HuR and STX5 act synergistically to load miRNAs into endosomes for their export. A cooperative miRNA binding by HuR and STX5 ensures the loading of miRNAs into endosomes. Interestingly, STX5 plays a dual role in miRNA export. The MVBs with miRNAs inside get fused to plasma membrane to release the content as EVs into extracellular space also in a STX5 dependent manner. Thus, STX5 not only ensures binding and endosome loading of miRNA, but also it facilitates membrane fusion in mammalian macrophages cells to ensure EV release. In combination, STX5 ensures selective miRNA export process in activated macrophage cells.

## Results

### Synchronised expression of Syntaxin 5 with miRNA-activity change in activated macrophage cells

Lipopolysaccharides (LPS), molecules derived from the outer membrane of Gram-negative bacteria, stimulate immune response in the macrophage cells by targeting Toll-like receptor 4 (TLR4) to activate NF-kB pathway^26^. During the early phase of LPS stimulation miRNPs get deactivated due to uncoupling of phosphorylated Ago2 and miRNAs that results in an enhanced expression of pro-inflammatory cytokines^9^. In this study, we have used LPS stimulated C6 rat glioblastoma cells to check the changes in miRNA activity and cellular levels after early and prolonged LPS exposure. We have found that the activity of let-7a was downregulated after early (4h) exposure but on prolonged exposure (24h) the activity was restored back to the normal. However, the cellular let-7a level remains largely unchanged throughout the LPS stimulation (Fig. 1A-C). We have also checked the change in the level of key factors of miRNA machinery, Ago2 and HuR, during the time dependent LPS stimulation in activated macrophages. Ago2 and HuR (Fig. 1D) are known to regulate the activity and export of miRNAs ^13^.We have also checked the level of one SNARE protein, Syntaxin 5 (STX5) that may play a crucial role in the exosomal export process in *C. elegans ^25^*. We have found a gradual increase in the STX5 level till 4hrs post LPS treatment and the level showed a steady decrease at later time points when miRNA activity was also restored (Fig. 1D-E).

**Figure 1.**
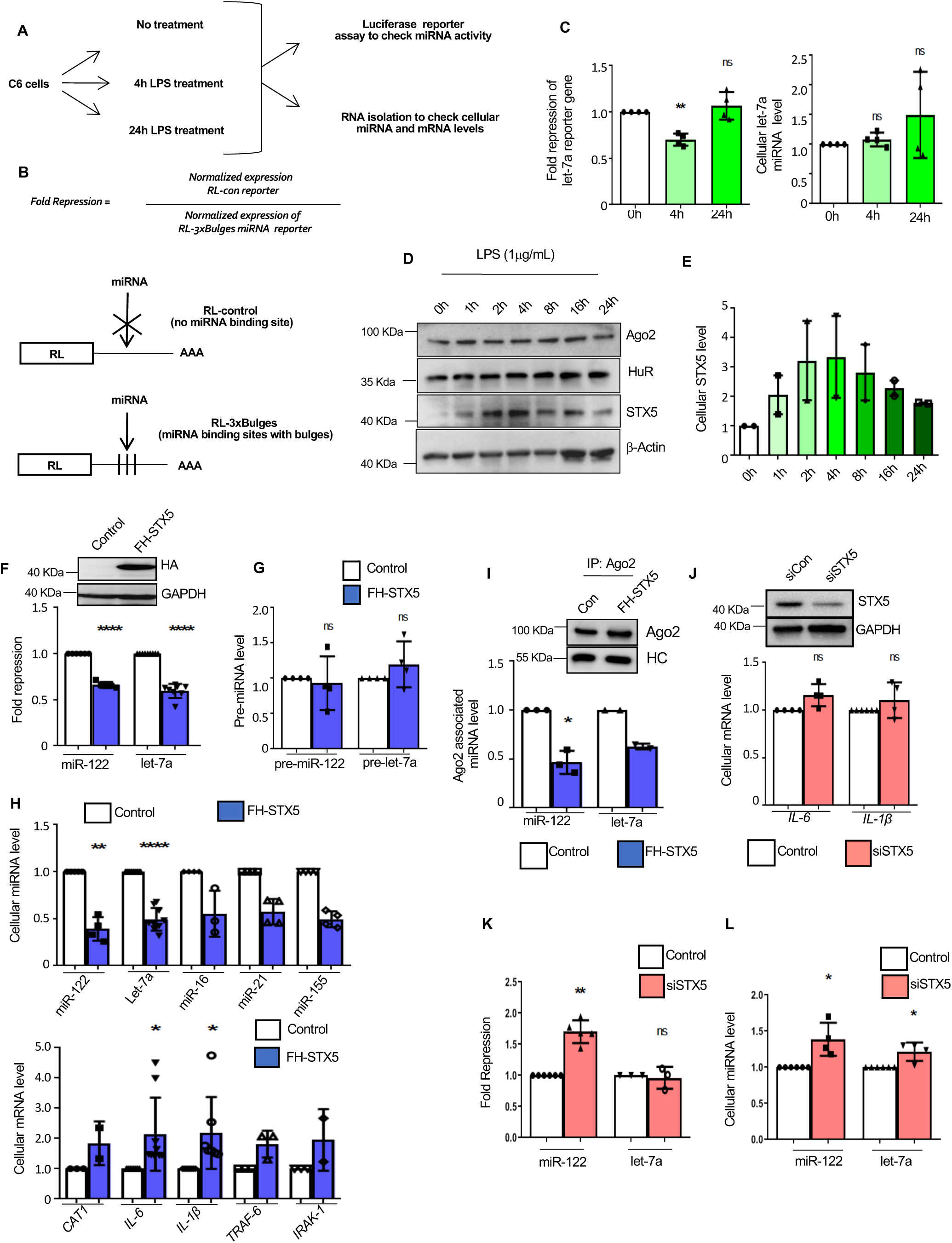
STX5 regulates cellular miRNA activity. (**A**) The effect of lipopolysaccharide, LPS (1μg/mL) on C6 glioblastoma cells was checked at different time points (0, 4 and 24hrs post LPS treatment). Cellular miRNA activity and endogenous miRNA level was measured. (**B)** Schematic outline of different Renilla luciferase (RL) reporter system used to evaluate miRNA activity. Calculation of fold repression is based on measurement of firefly luciferase normalized renilla luciferase vaules from control and experimental sets. (**C**) Cellular activity of let-7a was measured in control and LPS treated C6 cells by Renilla Luciferase assay (C, left panel, n=4 independent experiments; P= 0.0028, .04332), also the change in the total cellular let-7a level was quantified by qRT-PCR and normalized against U6 RNA level (C, right panel, n=4 independent experiments; P=0.249, 0.2655). (**D**) C6 cells were treated with LPS (1μg/mL) and harvested after 0hr, 1hr, 2hrs, 4hrs, 8hrs,16hrs and 24hrs post treatment. Western blot showed cellular levels of different proteins such as Ago2, HuR and STX5. Beta-Actin was used as the loading control. (**E**) Quantification was done and normalized (against beta-Actin) level of STX5 was graphically represented (n=2 independent experiments). (**F**) Cellular miR-122 and let-7a activity were measured C6 glioblastoma cells transfected with miR-122 expression plasmid (pmiR-122) and FH-STX5 or pCI-neo and renilla reporter plasmids for respective miRNAs (F, bottom panel, n≥5 independent experiments; P˂0.0001, ˂0.0001). Expression of FH-STX5 was also shown by Western blot analysis (F, top panel). (**G**)Relative cellular levels of pre-miR122 and pre-let-7a were also measured where GAPDH mRNA level was used as the normalizing control (n=4 independent experiments; P= 0.7394, 0.3089). (**H**) Cellular levels of different miRNAs were quantified in C6 cells expression using pCIneo or FH-STX5 expression plasmids. The miRNA levels were quantified by qRT-PCR and normalized against U6 snRNA level. (H, top panel, n≥3 independent experiments; P= 0.0023, ˂0.0001). Along with miRNAs, miRNA target mRNA levels such as CAT1, IL-6, IL-1β, TRAF-6 and IRAK-1 were also measured using qRT-PCR and normalized against GAPDH level (H, bottom panel, n≥2 independent experiments. For IL-6 and IL-1β, n≥7 independent experiments; P= 0.0218, 0.0389). (**I**) Immunoprecipitation (IP) of endogenous Ago2 from both control (pCI-neo) and FH-STX5 expressing C6 cells also expressing miR-122 and FH-Ago2 were performed followed by qRT-PCR based quantification of associated miR-122 and let-7a levels that were further normalized against the relative pulled down Ago2 levels (I, bottom panel, n≥2 independent experiments; P=0.016). Western Blot analysis was done to show the pulled down Ago2 level in the two sets along with the Heavy chain (I, top panel). (**J**) Endogenous STX5 level was knocked-down using siRNA against it in C6 cells and then mRNA levels of IL-6 and IL-1β were quantified and normalized against GAPDH levels (J, bottom panel, n=4 independent experiments; P=0.0742, 0.3431). The knock-down of STX5 protein level using siSTX5 in C6 cells was observed in the Western Blot analysis. GAPDH level was taken as the loading control (J, top panel). (**K**) siSTX5 transfected C6 cells were transfected with pmiR-122 and total miRNA activity for miR-122 and let-7a were measured using luciferase assay (n≥3 independent experiments; P= 0.0010, 0.7457). (**L**) In siSTX5 and pmiR-122 transfected C6 cells qRT-PCR data showing the cellular miR-122 and let-7a levels normalized against U6 RNA levels (n=4 independent experiments; P= 0.0425, 0.0439). In all experimental data, error bars are represented as mean ± SD. For statistical analysis, all experiments were done minimum three timesand P-values were calculated by two-tailed paired t test in most of the experiments unless mentioned otherwise; ns, non-significant, *P < 0.05, **P < 0.01, ***P < 0.001, ****P < 0.0001, respectively. Positions of molecular weight markers are marked and shown with the respective Western blots.

### Syntaxin 5 is necessary and sufficient for miRNA activity modulation in activated macrophage cells

We wanted to know whether this SNARE protein has a role to play in regulating the miRNA activity and their target mRNA expression. To check that we have transfected a FLAG-HA tagged version of STX5 (FH-STX5) in C6 cells and after 48h of transfection the miRNA activity was measured by Renilla Luciferase based reporter assay system for miRNA let-7a and miR-122. FH-STX5 expressing cells showed a significant decrease in the activity of both miR-122 and let-7a as compared to the control cells (Fig. 1F). qPCR data showed that along with the activity, cellular levels of different miRNAs were downregulated due to the expression of FH-STX5 but this decrease in the mature miRNA levels was not reflected in the levels of the precursor-miRNAs (Fig. 1G-H).Due to this reduction in the miRNA activity, respective target mRNA levels showed a significant upregulation (Fig. 1H). Downregulation of miRNA activity could be contributed by either lowering of miRNA levels or degradation of miRNP components or loss of miRNA-Ago2 interaction. To test the possibilities, we have expressed FH-STX5 in C6 cells and Ago2 immunoprecipitation was done using endogenous Ago2 specific antibody. A reduction in the association between Ago2 and miR-122 or let-7a was observed due to FH-STX5 expression as compared to the control cells but no change was observed in the Ago2 level (Fig. 1I).

To further confirm the role of STX5 in activity regulation of miRNAs, we have knocked-down the level of STX5 using an ON-target plus siRNA smart pool against it and measured the activity as well as the cellular miRNA/mRNA levels after 72h of siRNA transfection. We have found that the target cytokine levels such as IL-6 and IL-1β showed no change with respect to the siControl level (Fig. 1J). Cellular miR-122 activity was upregulated whereas let-7a activity showed no such change due to the knock-down of STX5 (Fig. 1K). However, the cellular miRNA levels of miR-122 and let-7a showed an increase in siSTX5-treated set respect to the siControl group (Fig. 1K) suggesting that knocking-down of STX5 has an opposite effect on cellular miRNA level compared to FH-STX5 expression. Overall, these results suggest that STX5 downregulates cellular miRNA activity by lowering the miRNA levels and by reducing the Ago2-miRNA interaction. Due to this reduced miRNA activity, cellular target mRNA and cytokine levels showed an enhanced expression in cells expressing FH-STX5.

Does reversible miRNA activity alteration during LPS treatment is controlled by STX5? To investigate that we transfected FH-STX5 or siSTX5 in C6 cells and treated the cells with LPS for 4h before harvesting and measured the miRNA activity and cellular levels of miRNA along with target mRNA levels. With respect to pCIneo control plasmid transfected cells, FH-STX5 expressing cells showed enhanced reduction in let-7a activity while the activity changes observed in siSTX5 treated cells was marginal (Fig. 2A-B). Interestingly, the reduced miRNA activity in cells expressing FH-STX5 could be contributed by reduced levels of let-7a miRNA whereas STX5 knocked-down cells showed an enhanced cellular expression of let-7a as compared to the control set (Fig. 2C). We have also checked the level of two pro-inflammatory cytokine mRNA levels IL-6 and IL-1β under the same conditions. As reported earlier, LPS treatment was found to induce the cellular cytokine expression. However, FH-STX5 showed a positive effect on induction of cytokine mRNA expression but knock-down of STX5 resulted in a significant drop in the levels of both the cytokine mRNA as compared to the control cells (Fig. 2D).

**Figure 2.**
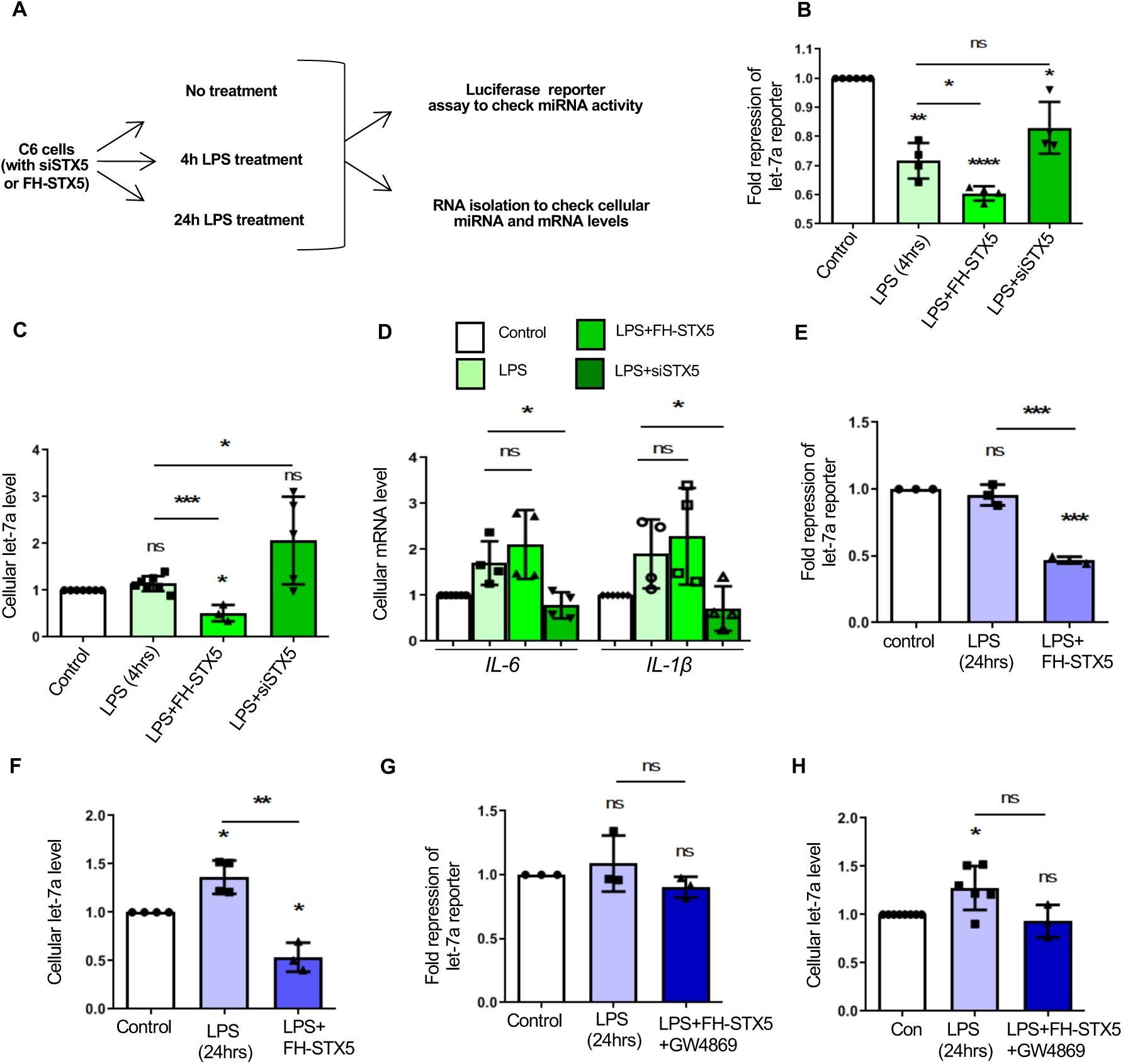
STX5 is necessary for LPS mediated derepression of miRNA activity in mammalian macrophages. (**A**) The effect of LPS (1μg/mL) on C6 glioblastoma cells was checked at different time points (0, 4 and 24hrs post LPS treatment). Cellular let-7a activity and endogenous let-7a level was measured. (**B-D**) In addition to the 0 and 4hrs of LPS treatment, C6 cells were also transfected with FH-STX5/siSTX5 and change in the cellular let-7a activity was measured by Luciferase assays (B, n=4 independent experiments; P=0.0027, <0.0001, 0.0315; unpaired t-test, P=0.0142, 0.0816). The change in the total cellular let-7a level was quantified against U6 level (C, n≥3 independent experiments; P= 0.0592, 0.0402, 0.0644; unpaired t-test, P=0.0005, 0.0265). qRT-PCR was done to check the cellular target cytokine levels. *IL-6* and *IL-1β* levels were measured and normalized against GAPDH level. (D, n=4 independent experiments; unpaired t-test, P=0.3970, 0.0161, 0.5748, 0.0376) (**E,F**) Both control and FH-STX5 expressing C6 cells were treated with 1μg/mL LPS for 24hrs. (**E**)Total cellular let-7a activity was measured for each set using Luciferase based assay system (n=3 independent experiments; P=0.4289, 0.0007; unpaired t-test, P=0.0005). (**F**) qRT-PCR showing the total cellular let-7a levels normalized with U6 (n≥3 independent experiments; P= 0.0246, 0.0332; unpaired t-test, P=0.0012). (**G,H**) Both control and FH-STX5 expressing C6 cells were treated with 1μg/mL LPS for 24hrs. GW4869 was added to the system and incubated for 20hrs. DMSO was added in the control sets. Cellular let-7a activity was measured and analyzed (G, n=3 independent experiments; P=0.555, 0.1746; unpaired t-test, P=0.2410) and cellular level of let-7a was also quantified and normalized against U6 RNA level (H, n≥3 independent experiments; P=0.0310, 0.5550; unpaired t-test, P=0.0553) In all experimental data, error bars are represented as mean ± SD. For statistical analysis, all experiments were done minimum three times and P-values were calculated by two-tailed paired t test in most of the experiments unless mentioned otherwise; ns, non-significant, *P < 0.05, **P < 0.01, ***P < 0.001, ****P < 0.0001, respectively. Positions of molecular weight markers are marked and shown with the respective Western blots.

Restoration of miRNA activity after prolonged LPS exposure has been reported earlier^9^. Interestingly, upon 24h of LPS treatment, a significant downregulation of let-7a activity was found upon FHA-STX5 expression compared to non-expressing control group (Fig. 2E). This effect was also reflected in the cellular miRNA level and FH-STX5 expressing cells showed a significant reduction in the let-7a level as well (Fig. 2F). Altogether these findings indicate that STX5 mediated lowering of miRNA activity in LPS stimulated cells happened due to miRNA lowering.

Lowering of total cellular miRNAs can happen due to various reasons like extracellular export via extracellular vesicles or due to miRNA degradation or due to retarded miRNA transcription/processing steps. As no change was observed in the precursor forms of miRNAs on STX5 level manipulation (as showed in Fig. 1G), the possibility of faulty transcription/processing may not be the primary reason of observed changes in mature miRNA levels. Next, we wanted to check whether the export pathway plays any role in this STX5 mediated lowering of miRNAs. Extracellular Vesicles or Exosomes play an important role in the export of different molecules such as miRNAs, mRNAs or proteins from various types of mammalian cells including macrophages ^27^. GW4869, a neutral sphingomyelinase (nSMase) inhibitor, is the most widely used pharmacological agent for blocking exosome biogenesis^28^. We have expressed FH-STX5 in C6 cells and treated the cells with LPS along with GW4869 before analysing miRNA activity and level check. In cells, where perturbation of export pathway by GW4869 treatment was done, FH-STX5 failed to downregulate both the cellular activity and level of let-7a (Fig. 2G-H). This data indicates the importance of extracellular export in the STX5 mediated lowering of miRNA activity.

### Syntaxin5-mediated miRNA activity lowering prevents infection of mammalian macrophages by invading pathogen *Leishmania donovani* both *ex vivo* and *in vivo*

*Leishmania donovani*(*Ld*), the causative agent of visceral leishmaniasis infects the host macrophage cells and escapes the immune response by inhibiting ROS generation, pro-inflammatory cytokine production and by inducing the expression of anti-inflammatory cytokine IL-10 ^29^. From our observations, it is obvious that STX5 mediated lowering of miRNA activity results in an enhanced expression of pro-inflammatory cytokine mRNAs inside the macrophage cells. We were curious to know whether this pro-inflammatory cytokine surge has any role to play in regulating the host pathogen interaction in the macrophage cells. To test that we have expressed FH-STX5 in murine macrophage cell RAW264.7, and after 48h of transfection the cells were infected with *Leishmania* parasite and incubated for 24hbefore harvesting. Along with *Ld*, GW4869 was also added to one experimental set to check whether exosomal export pathway control by STX5 may affect the infection event. As reported earlier ^30^, *Ld* lowers the miRNA activity while the relative cellular miRNA level goes up in infected macrophages (Fig. 3A-B). FH-STX5 expressing RAW264.7 cells, infected with *Ld*, showed downregulation of both the activity and cellular level of let-7a. Interestingly, GW4869 treated FH-STX5 expressing infected cells showed no or little change in let-7a level and activity as compared to the pCIneo control plasmid transfected and *Ld* infected cells (Fig. 3A-B). Endogenous Ago2 was pulled down from the control and FH-STX5 expressing infected RAW264.7 cells. Interaction between Ago2 and let-7a was reduced due to FH-STX5 expression, but the total pulled-down Ago2 level did not change (Fig. 3C).

**Figure 3.**
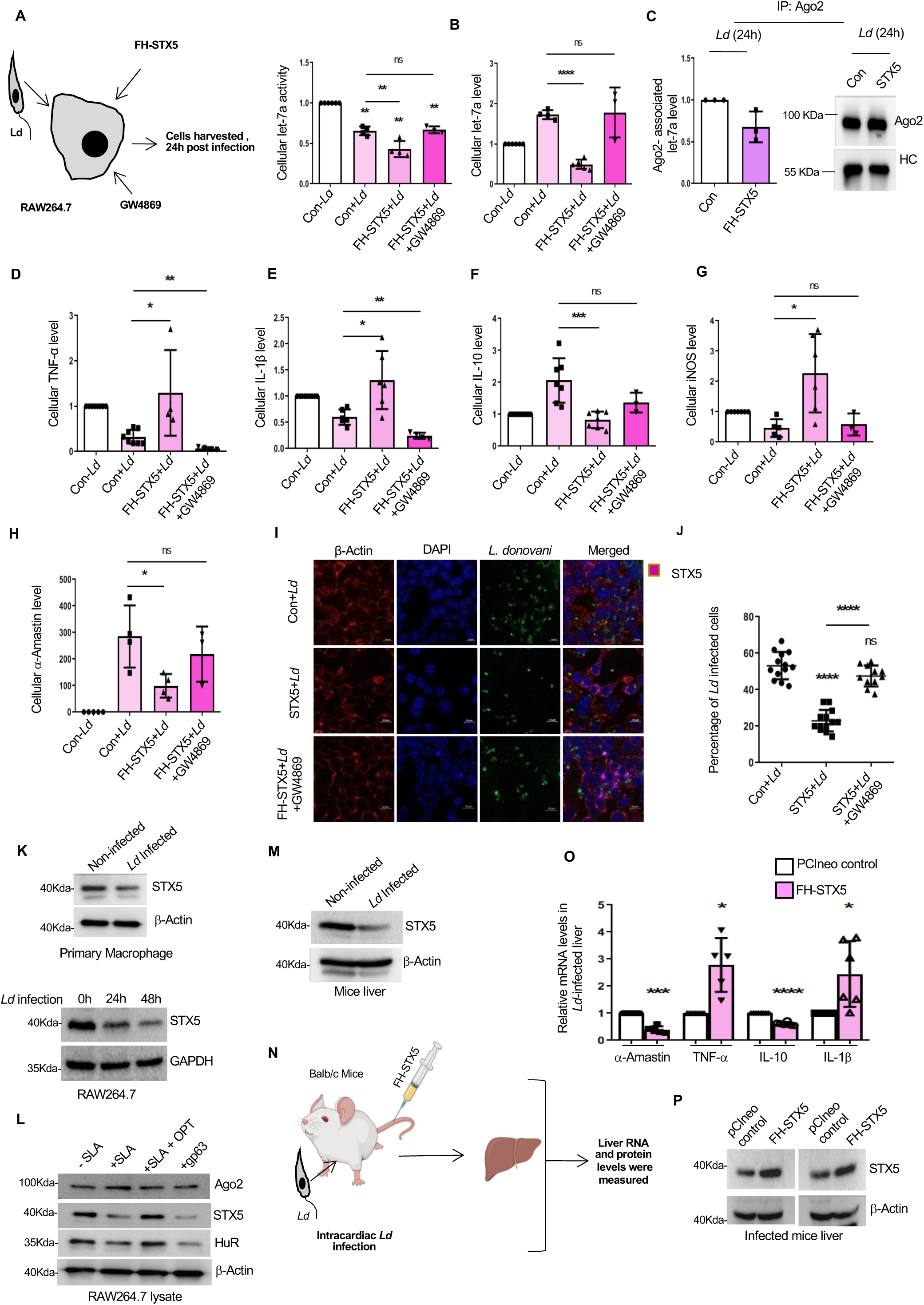
STX5 clears internalized pathogens by inducing expression of inflammatory cytokines in mammalian macrophages. (**A-B**) RAW264.7 cells expressing FH-STX5 were infected with *Leishmania donovani* strain Ag83 at a ratio of 1:10 (macrophage:*Ld*). Along with this *Ld* infection, one set of cells was also treated with GW4869 and incubated for 24hrs before harvesting. Luciferase based assay system was used to measure the relative let-7a activity for each set (A, n≥3 independent experiments; P=0.0011, 0.0015, 0.0053; unpaired t-test, P=0.0078, 0.7374). (**B**) qRT-PCR was done to check the cellular let-7a level change in each set. U6 was taken as the normalizing control. (n≥3 independent experiments; unpaired t-test, P< 0.0001, 0.8701). (**C**) Immunoprecipitation was done with anti-Ago2 antibody from *Ld* infected RAW264.7 cells expressing FH-STX5 or pCIneo. Ago2 associated let-7a level was checked by qRT-PCR. (C, left panel, n=3 independent experiments) and pulled-down Ago2 level along with the heavy chain was also shown (C, right panel). (**D-H**) In the above-mentioned experimental sets (in A,B panels), qRT-PCR was done to check different target cytokine and other mRNA levels such as TNF-α (D, n≥4 independent experiments; unpaired t-test, P=0.0141, 0.0086), IL-1β (E, n≥4 independent experiments; unpaired t-test, P=0.0130, 0.0018), IL-10 (F, n≥3 independent experiments; unpaired t-test, P=0.0009, 0.1428), iNOS (G, n≥3 independent experiments; unpaired t-test, P=0.0140, 0.6269) and α-Amastin level (H, n≥3 independent experiments; unpaired t-test, P=0.0249, 0.4726). α-Amastin level denotes the amastigote stage of parasite population inside the infected cells and a representation of infection level. GAPDH was used as the normalizing control. (**I-J**) Similarly, RAW264.7 cells expressing FH-STX5 were infected with CFSE stained *Ld* at a ratio of 1:10 (macrophage:*Ld*). GW4869 was added to on each set of infected cells around the same time i.e. 24hrs before harvesting. Confocal microscopy was done to check the of internalized *Ld* number in infected RAW264.7 cells expressing or not expressing FH-STX5 (I). Beta-actin was used to mark the cellular boundary (red), CFSE stained *Ld* was shown in green and FH-STX5 expressing are shown in purple. Nuclei were stained with DAPI (blue). Percentage of *Ld* infection or internalization was calculated and graphically shown (J, n≥15 independent field of views were analyzed; unpaired t-test, P= <0.0001, 0.0551, <0.0001).(**K**) Mouse primary peritoneal macrophage cells and RAW264.7 cells were infected with *Ld*. Western blot was done to check the endogenous level of STX5 in both the *Ld* infected and non-infected primary macrophage cells (K, top panel) and RAW264.7 cells (K, bottom panel). Beta-actin was taken as the loading control for primary macrophage and GAPDH was taken as the loading control for RAW264.7 cells. (**L**) RAW264.7 cell lysate was incubated with soluble *Leishmanial* antigen (SLA), isolated from *Leishmania donovani* lysate, for 30mins at 37°C. Another set of RAW264.7 cellular lysate was treated with SLA which was pre-incubated with 10μM *o*-phenanthroline for 30mins at 4°C. RAW264.7 lysate was also incubated with *Leishmania* derived purified metalloprotease, gp63 for 30mins at 37°C. Western blot was done to check the endogenous level of cellular Ago2, HuR and STX5 for all the above-mentioned sets. Beta-actin was used as the loading control. (**M**) Western blot analysis was done to check the endogenous level of STX5 in *Ld* infected and non-infected mice liver samples (infected for 30 days). Beta-actin was taken as the loading control. (**N**) Schematic of the animal experiment where 4-6weeks old male Balb/c mice were injected with pCI-neo or F.HA-STX5 plasmid via tail vein and intracardiacally infected with *Ld* for 35days before sacrificing. Liver samples were collected and further processed for subsequent RNA and protein expression check. (**O**) RNA was isolated from all the liver samples and different target cytokine and other mRNA levels such as α-Amastin (to measure the level of *Ld* infection), TNF-α, IL-10 and IL-1β were checked and normalized against the GAPDH level. (Biological replicates=3, technical replicates≥5; P=0.0004, 0.0161, <0.0001, 0.0331). (**P**) Western blot showed the level of FH-STX5 in both control and FH-STX5 encoding plasmid injected mice liver tissues. Beta-actin was taken as the loading control. In all experimental data, error bars are represented as mean ± SD. For statistical analysis, all experiments were done minimum three times and P-values were calculated by two-tailed paired t test in most of the experiments unless mentioned otherwise; ns, non-significant, *P < 0.05, **P < 0.01, ***P < 0.001, ****P < 0.0001, respectively. Positions of molecular weight markers are marked and shown with the respective Western blots.

Next, we focused on the level of cytokine and other target mRNAs in host macrophages under infection conditions. We have checked different target mRNA levels such as TNF-α, IL-1β, IL-10, iNOS and α-Amastin. As expected,*Ld* infected cells showed a downregulation in the pro-inflammatory cytokine levels (TNF-α, IL-1β) and an enhanced expression of anti-inflammatory IL-10 (Fig. 3D-F). On the other hand, FH-STX5 expressing infected cells showed an overall pro-inflammatory cytokine response in macrophage cells by promoting the production of TNF-α, IL-1β and by inhibiting anti-inflammatory IL-10 production (Fig. 3D-F). iNOS (inducible nitric oxide synthase) is an enzyme that produces nitric oxide (NO) which plays an important role in killing the internalized *Leishmania* parasite by the host macrophages ^31^. Therefore, greater the level of iNOS, higher is the level of NO production and consequently lesser is the chance of pathogen survival inside the host cells. Normal *Ld* infected cells showed a lesser expression level of iNOS whereas FH-STX5 expressing infected cells showed a significant upregulation of the same (Fig 3G). But interestingly, when the exosomal biogenesis was blocked with GW4869 in infected cells, FH-STX5 failed to induce any such pro-inflammatory response inside the macrophages which was evident from the downregulation of TNF-α, IL-1β and iNOS and simultaneous upregulation of IL-10 (Fig. 3D-G).

Now, to check the level of parasite internalization we have measured the cellular α-Amastin level that denotes the internalized *Leishmania* amastigote population in the host macrophage cells. The result showed that FH-STX5 was able to lower the amastigote specific amastin level as compared to PCIneo control transfected and *Ld* infected cells whereas GW4869 treatment reversed the lowering of amastin level due to FH-STX5 expression in infected RAW264.7 cells (Fig. 3H). Confocal microscopy analysis of the infected cells was also done to check the level of *Ld* internalization(Fig. 3I-J). The percentage of *Ld* infection was quantified, and it was observed that FH-STX5 was able to prevent the internalization of *Leishmania* into the host macrophage significantly as compared to the infected pCIneo control plasmid transfected RAW264.7 cells. However, the infection load again went back to a higher level due to the GW4869 treatment of infected cells even after expression of FH-STX5 (Fig. 3I-J). Therefore, it can be said that STX5 mediated lowering of miRNA activity and cellular level induces a pro-inflammatory response inside the *Ld* infected host murine macrophage cells both by promoting the pro-inflammatory and suppressing the anti-inflammatory cytokine expression that ultimately prevent the internalization of the invading pathogen significantly. But this STX5 mediated regulation of host-pathogen interaction gets reversed when the host cells were treated with GW4869 to block the exosomal pathway.

Can STX5 control the infection process in the *in vivo* context? Both in liver tissue or murine primary peritoneal exudates cells (PEC) or murine macrophage RAW264.7 cells, the endogenous level of STX5 showed a reduction due to *Ld* infection as compared to the non-infected control set (Fig. 3K). In RAW264.7 cells the endogenous STX5 level showed a further decrease at 48hrs post infection (Fig. 3K). If we incubate Soluble Leishmanial Lysate (SLA) with RAW264.7 cell extract, SLA targets endogenous STX5. We detected degradation of endogenous STX5 but this reduction in STX5 level was reversed when SLA was pre-incubated with the Zn^2+^ chelator ortho-phenanthroline (OPT) (Fig. 3L). It has been reported earlier that Leishmanial Zn-metalloproteases are involve in the cleavage of HuR and Dicer1 protein in infected cells to restrict miRNA activity. GP63 is the most abundant Zn-metalloprotease that cleave DICER1 and HuR in *Ld* infected cells ^13,32^. With purified GP63, we found significant lowering of STX5 level also along with HuR protein upon incubation of RAW264.7 cell extract. Beta actin level did not show much change (Fig. 3L). Hence it can be assumed that *Leishmanial* Zn-metalloprotease GP63 may cleave cellular STX5 protein to downregulate the pro-inflammatory cytokines that ensures better internalization of the pathogen inside the host macrophages. Level of STX5 also get downregulated in the liver of *Ld* infected mice (Figure 3M). *Leishmania* thus targets endogenous STX5 to downregulate for its survival. Can exogeneous expression of FH-STX5 reverse the infection process? To test, 4-6weeks male Balb/C mice were injected with pCIneo or FH-STX5 expression plasmid via tail-vein route followed by intracardiac *Leishmania* infection. After 35days of repeated injection of FH-STX5 expression plasmid at regular intervals, the mice were sacrificed, and liver tissue and blood samples were collected for further protein and RNA analysis (Fig. 3N). As liver is the first tissue that gets infected during *Ld* infection process, we isolated RNA from the homogenized liver tissue samples and target cytokine and cellular miRNA levels were measured. Like the *ex vivo* observations, we have found that in the infected liver, pro-inflammatory cytokines (TNF-α and IL-1β) showed a significant increase along with a reduction in the anti-inflammatory IL-10 level upon FH-STX5 expression (Fig. 3O). Moreover, α-Amastin level also showed a significant drop on FH-STX5 expression (Fig. 3O) suggesting that FH-STX5 mediated pro-inflammatory response in the liver tissue clears *Leishmania* infection in the infected mice liver. Total tissue miRNA levels in infected liver was also in alignment with the cell line experiments as most of the miRNAs like miR-122, let-7a or miR-155 showed a significant downregulation upon FHA-STX5 expression in the infected mice. However, miR-146a, an anti-inflammatory miRNAs known to get upregulated in Ld infected macrophage ^33^, don’t get reduced by FH-STX5 (Supplementary Figure S1A).

Other than liver tissue, miRNAs were also checked from the serum samples of the infected mice. Serum isolated from the peripheral blood also contains circulating miRNAs packaged in extracellular vesicles^34^. All three miRNAs that we have checked in serum samples such as miR-122, let-7a and miR-146a showed upregulation in mice expressing FH-STX5 (Supplementary Figure S1B-C). Western blot analysis using anti-STX5 antibody confirmed the enhanced expression of STX5 in mice injected with FH-STX5 plasmid (Fig. 3P). Altogether, excess expression of STX5 was able to prevent the *Leishmanial* infection in the liver tissues of infected mice by lowering the miRNA levels that may contributes to upregulating of pro-inflammatory cytokine expression.

### Syntaxin 5 regulates miRNA activity by promoting its export via EVs

As mentioned earlier that STX5 mediated lowering of miRNAs can happen due to various reasons such as faulty transcription/processing of precursor miRNAs, export via extracellular vesicles or by its degradation. No change in the cellular level of the precursor miRNAs suggested that the transcription process was not affected due to the FH-STX5 expression. Next to check the export pathway, we have expressed FH-STX5 in C6 cells and conditioned media or culture supernatant was collected to get the extracellular vehicles (EVs) secreted by respective cells (Fig. 4A). EVs were characterized by Nano-particle tracking analysis (Fig. 4B). Data obtained from the NTA showed that there was a significant increase in the number of EVs released from the FH-STX5 expressing cells as compared to the control cells (Fig. 4C). No such change was observed between the sizes of the EVs. Different protein factors were also checked and as expected Ago2 and HuR were not detected in the EV but the exosomal marker proteins such as Alix and Flotillin-1 showed an increase in their levels due to FH-STX5 expression which further strengthens our finding that FH-STX5 promotes exosomal export (Fig. 4D). Moreover, presence of FH-STX5 in EV fraction was detected in the Western blot suggesting that FH-STX5 not only promotes the EV export, but also gets packaged into the EVs as well (Fig. 4D). Next, we measured the miRNA content in the EVs and qPCR data normalized against the number of EVs released showed that the level of miR-122 and let-7a packaged per EV were significantly upregulated upon FH-STX5 expression (Fig. 5E). So STX5 facilitates both the EV number and the EV-associated miRNA levels that might explain the lowering of cellular miRNAs observed in presence of FH-STX5. To confirm that STX5 facilitates EV-export, we have knocked-down the expression of STX5 using siSTX5 and EVs were isolated from the conditioned media and analysed further by NTA (Fig. 4F). NTA data showed a significant drop in the EV number due to STX5 knock-down (Fig. 4G) suggesting that STX5 plays a crucial role in the genesis of the extracellular vesicles. However, the level of miRNAs (miR-122 and let-7a) that are getting exported out of the cells per exosome showed a non-significant change as compared to the control although total level of EV associated miRNA was decreased (Fig. 5H). Overall, when the STX5 level was knocked-down the total miRNAs that were getting exported out via EVs showed a reduction and the effect was also reflected in the increase in cellular miRNA levels.

**Figure 4.**
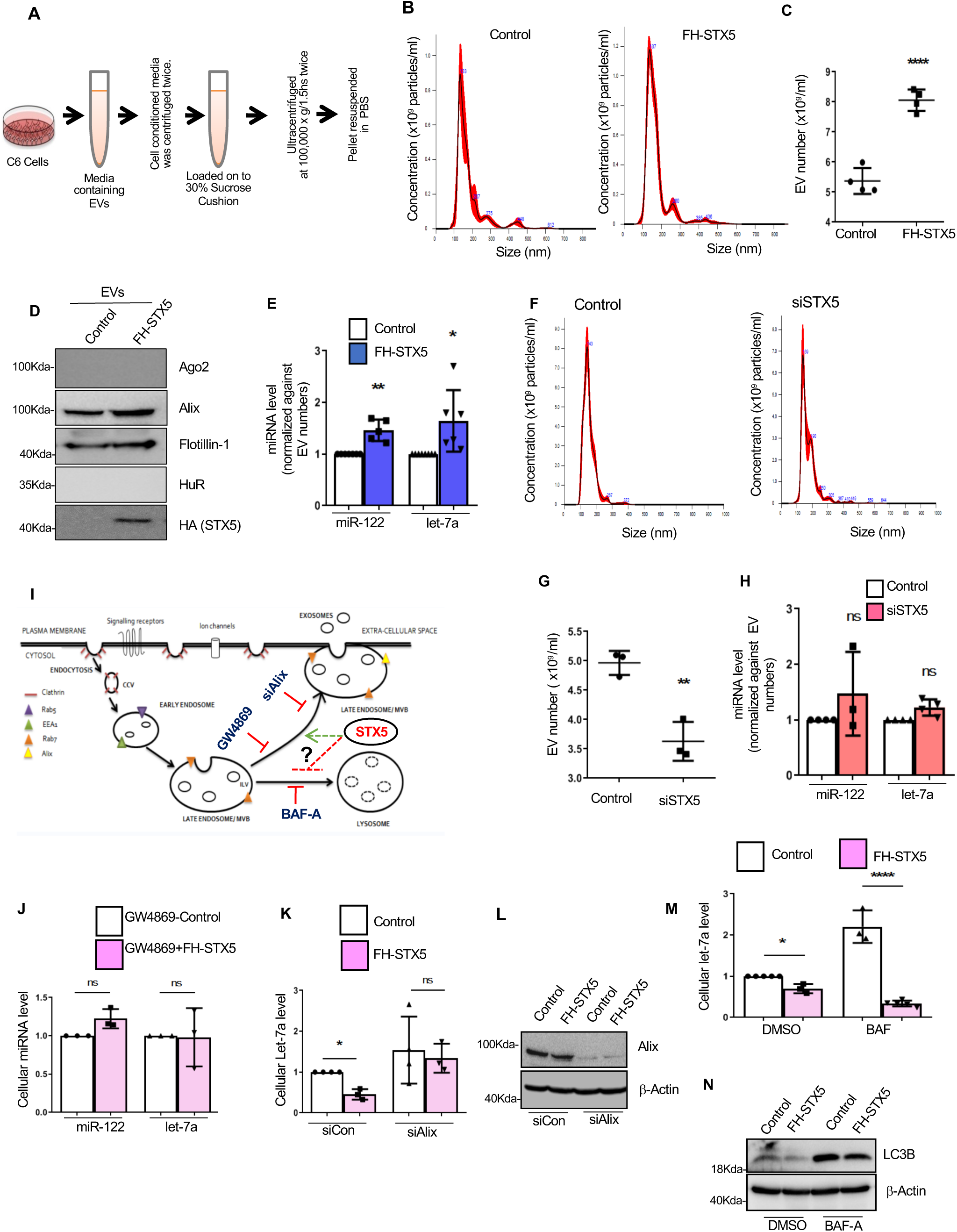
STX5 lowers cellular miRNAs by accelerating their export. (**A**) C6 cells expressing FH-STX5 or transfected with pCIneo (control) were allowed to grow in the exosome depleted media for 24hrs before harvesting. Exosomal vesicles were isolated from the conditioned media by ultracentrifugation. (**B**) Isolated extracellular vesicles were resuspended in PBS and subsequently analyzed and characterized by a Nano-particle tracking analyzer(NTA). (**C**) Number of EVs released from the both control and FH-STX5 expressing cells were quantified by NTA (n=4 independent experiments; unpaired t-test, P<0.0001). (**D**) Western blot analysis showed the level of different marker proteins such as Alix and Flotilin1 in the EV samples derived from control and FH-STX5 expressing cells. It also showed the presence of FH-STX5 in the EVs from FH-STX5 expressing cells. Ago2 and HuR were not detected in EV fractions. (**E**) qRT-PCR was done to check the miRNA content in EVs. Equal amount of RNA was used as the template to check the miR-122 and let-7a levels in the EV fraction and miRNA levels were normalized either against the number of EVs or against the level of Flotilin1 in the respective sets (n≥5 independent experiments; P=0.0077, 0.0459). (**F**) EVs were isolated from the conditioned media of both siControl and siSTX5 treated C6 cells using ultracentrifugation and those vesicles were subsequently characterized by Nano-particle tracking analyzer. (**G**) Number of EVs released from both these sets were quantified by NTA and showed in the graph (n=3 independent experiments; unpaired t-test, P=0.0040). (**H**) miR-122 and let-7a levels were checked for both siControl and siSTX5 treated cells. Equal amount of RNA was used for the qRT-PCR reaction and the miRNA levels were normalized against the number of EVs released from each set (n=3 independent experiments; P=0.3929, 0.1742). (**I**) Schematic diagram to show the endocytic pathway and the steps which were perturbed to stop the fusion of late endosomal vesicles to either plasma membrane or lysosomes by adding different chemical compounds or siRNA to check the effect of STX5 on late endosomal fusion events. (**J**) C6 cells were transfected with pCI-neo (control) or FH-STX5 expression plasmids and cells from both the sets were treated with 10μM GW4869, 24hrs before harvesting. Cellular miR-122 and let-7a levels were measured against the U6 content (n=3 independent experiments; P=0.0912, 0.9417). (**K**) C6 cells were transfected with pCIneo (control) and FH-STX5 encoding plasmids and cells from both the sets were treated with siControl and siAlix for 72hrs. qRT-PCR was done to check the change in the cellular let-7a level against U6 (n≥3 independent experiments; P=0.0183; unpaired t-test, P=0.7165). (**L**) Change in the cellular Alix level due to siRNA treatment was shown in the Western blot image, beta-actin level was considered as the loading control. (**M**) To perturb the lysosomal targeting of late endosomes, both control and FH-STX5 expressing cells were treated with bafilomycin-A (BAF-A) or DMSO for 8hrs. Cellular let-7a level was measured and normalized against U6 level (n≥3 independent experiments; P=0.0441; unpaired t-test P= <0.0001). (**N**) Western blot image showed a significant change in the autophagic marker LC3B level due to bafilomycin treatment as compared to DMSO control. Beta-actin was taken as the loading control. In all experimental data, error bars are represented as mean ± SD. For statistical analysis, all experiments were done minimum three times and P-values were calculated by two-tailed paired t test in most of the experiments unless mentioned otherwise; ns, non-significant, *P < 0.05, **P < 0.01, ***P < 0.001, ****P < 0.0001, respectively. Positions of molecular weight markers are marked and shown with the respective Western blots.

**Figure 5.**
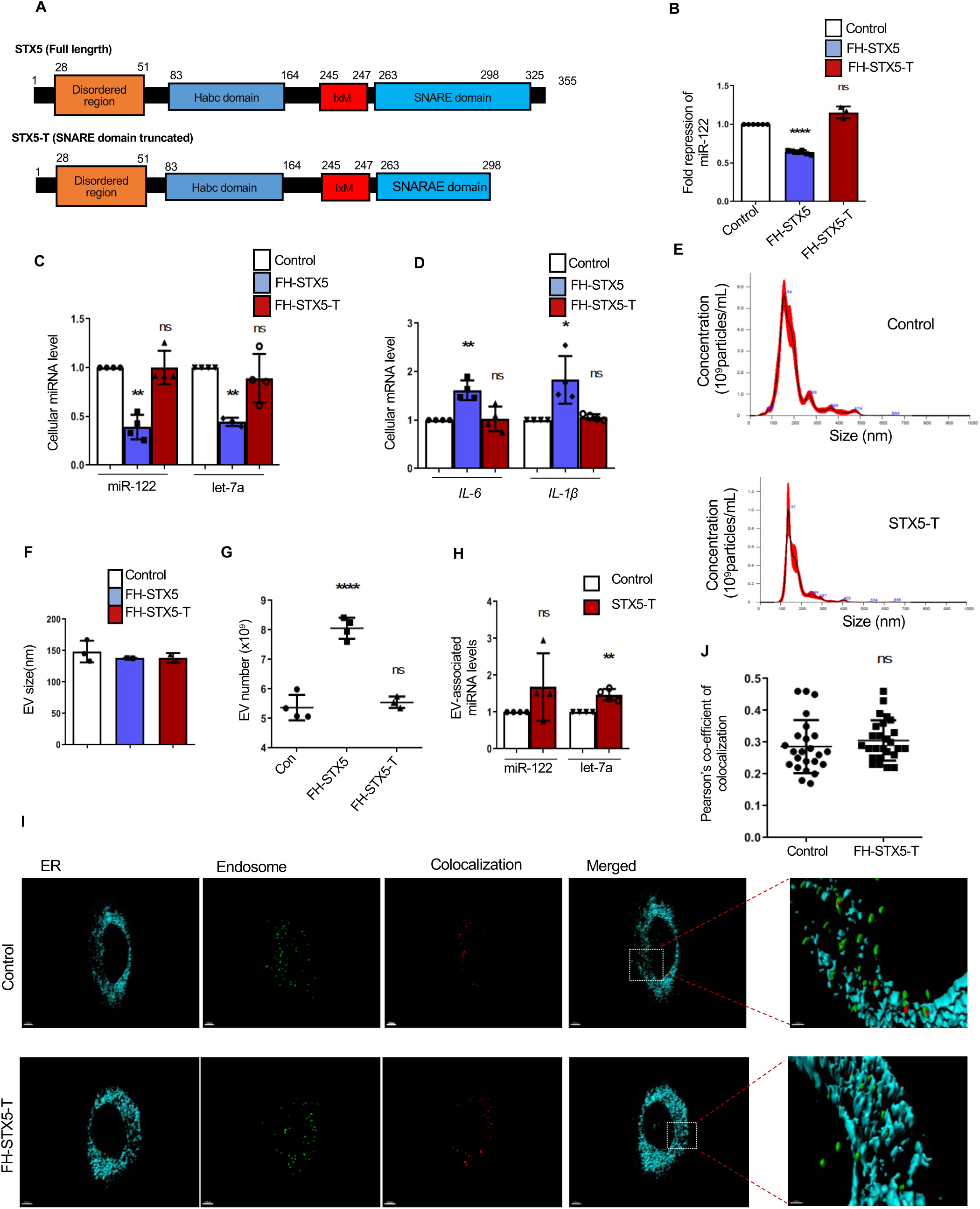
The SNARE domain of STX5 is necessary for miRNA export function. (**A**) The map of full length STX5 and truncated STX5 (STX5-T) were shown along with their structural domains. A truncation in the full length STX5 was made by deleting a stretch of 57 amino acids from the C-terminal SNARE domain and this truncated protein was expressed in C6 cells. (**B**) Luciferase based assay system was used to check the cellular miR-122 activity in FH-STX5 and FH-STX5-T expressing C6 cells (n≥3 independent experiments; P= <0.0001, 0.0735). (**C-D**) qRT-PCR data showed the change in both the cellular miRNA and cytokine levels upon expression of FH-STX5 or FH-STX5-T expression. miR-122 and let-7a levels were measured and normalized against U6 snRNA level (C, n≥3 independent experiments; P=0.0023, 0.9987, 0.0019, 0.4460) and IL-6 and IL-1b levels were measured and normalized against cellular GAPDH level (D, n=4 independent experiments; P=0.0091, 0.8492, 0.0428, 0.1290). (**E**) EVs were isolated from the conditioned media of pCIneo (control) or FH-STX5 or FH-STX5-T expressing cells using ultracentrifugation and subsequently characterized by Nano-particle tracking analyzer. (**F-G**) Nano-particle tracker data showed the size (**F**) and also the number of the EVs isolated from the cells in the each set of experiments (**G,** n≥3 independent experiments, unpaired t-test; P=<0.0001, 0.5431). (**H**) qRT-PCR data showed the miR-122 and let-7a content in EVs isolated from both control and FH-STX5-T expressing cells. Equal amount of RNA from each experimental set were used for the assay and the miRNA levels were normalized against the number of EVs released from the respective sets(n=4 independent experiments; P=0.2340, 0.0090). (**I**) Confocal imaging was done to check the effect of FH-STX5-T expression on the interaction between endoplasmic reticulum (ER) and early endosomal vesicles (EE) in C6 cells where ER was tagged with ER-dsRed plasmid (cyan), early endosmoes were detected with anti-EEA1 antibody via indirect fluorescence(green) and colocalized regions were shown in red. Enlarged colocalized regions/bodies were also shown in right most panel. (**J**) Pearson’s co-efficient of colocalization was calculated for both control and FH-STX5-T expressing cells and represented graphically (n≥24 cells from different field were analyzed; unpaired t-test, P=0.3648). In all experimental data, error bars are represented as mean ± SD. For statistical analysis, all experiments were done minimum three times and P-values were calculated by two-tailed paired t test in most of the experiments unless mentioned otherwise; ns, non-significant, *P < 0.05, **P < 0.01, ***P < 0.001, ****P < 0.0001, respectively. Positions of molecular weight markers are marked and shown with the respective Western blots.

EVs are primarily released from the mammalian cells when the multi-vesicular bodies (MVBs) fuse with the plasma membrane to release their interlunimal vesicles (ILVs) into the extracellular milieu as “exosomes”^35^. MVBs can either go fuse with the plasma membrane to release EVs or it can fuse with the lysosomes for degradation. Now previous results showed that STX5 promotes the EV-mediated export. But how it affects the endosomal/MVBs trafficking to lysozomes? To check how it affects the intracellular trafficking of MVBs, we have perturbed the fusion of MVBs to plasma membrane or lysozomes individually. We have mentioned earlier that GW4869 blocks ceramide dependent exosome biogenesis by inhibiting neutral sphingomyelinase (nSMase) ^28^. Alix is an ESCRT-III associated protein that plays a key role in the fusion process of MVBs to the plasma membrane to release exosomes in exracellular space and hence knocking-down of Alix should result in lowering of EV-release from the cells ^36^. Alternatively, lysosomal targeting of MVBs can be blocked by Bafilomycin A (BAF-A), a macrolide that targets V-ATPase to prevent the acidification of the endosome essential for its fusion with lysozomes^37^.FH-STX5 expressing cells showed an enhanced EV-export but whether it has any effect on the lysosomal targeting of MVBs was an unexplored question. We have measured the cellular miRNA levels from each set of experiments to check what part of the endocytic pathway is responsible for STX5 mediated lowering of cellular miRNAs. Interestingly, when the cells were treated with GW4869, STX5 was unable to lower the cellular miR-122 and let-7a levels (Fig. 4I-J). Similar findings were also observed when the cells were transfected with siAlix. In the siControl set,FH-STX5 showed significant lowering of let-7a level but when Alix was knocked-down, no such difference was found in the let-7a level between control and FH-STX5 expressing cells (Fig. 4K-L). But interestingly, after treatment of BAF-A (DMSO was used in the control set), FH-STX5 was able to lower the cellular let-7a level (Fig. 4M). As BAF-A is a known inhibitor of autophagy, the autophagic marker protein LC3B showed an upregulation in BAF-A treated cells as compared to the DMSO treated cells (Fig. 4N). Overall, these data suggest that STX5 mediated lowering of cellular miRNAs is primarily dependent on the enhanced EV-release, but it does not get affected if the lysosomal targeting of MVB is perturbed.

### Syntaxin 5 promotes EV-mediate miRNA export by enhancing its accumulation in MVBs and prevents targeting of miRNA-loaded MVBs to lysozomes

It has been reported earlier that miRNA activity heavily depends on the endocytic pathway and miRNA mediated repression and target mRNA degradation are two spatio-temporally uncoupled events ^4^. In this study, we have seen STX5 promotes the extracellular export of miRNA but we wanted to check what effect does it have on the sub-cellular localization of miRNAs. To do so, differential centrifugation was done to separate the different cellular organelles based on their density on an Optiprep^TM^ Gradient and fractions were collected and used for further protein and RNA isolation and analysis (Fig. S2A). Early fractions were positive for Endosomal proteins EEA1, Alix and HRS positive and negative for ER protein Calnexin, whereas the late fractions enriched for ER, were Calnexin positive but EEA1, Alix and HRS negative. STX5 was found to be present in both the endosomal and ER fractions (Fig. S1A). RNA was isolated from both the endosomal and ER fraction and qPCR data showed that miR-122 and let-7a were getting more enriched in the endosomal compartment as compared to the ER due to FH-STX5 expression (Fig. S2B-C). Total polysomal fraction was isolated to check the associated miRNA levels in control and FH-STX5 expressing C6 cells (Fig. S2D). Polysomal fraction was positive for marker protein L7a (Fig. S2D). RNA samples were isolated from the polysomal fraction and analysed further to check the associated miRNA levels. FH-STX5 expressing cells showed lesser enrichment of miR-122 and let-7a in the polysomes (Fig. S2E) which was consistent with the sub-cellular fractionation data, suggesting miRNA accumulation in endosomal fraction in cells expressing FH-STX5.

Next to check the effect of STX5 on lysosomal targeting of the miRNA loaded MVBs, we did confocal imaging to quantitatively measure the level of interaction between late endosomes/MVBs (marked with Rab7, green) and lysosomes (marked with lysotracker, red) in both FH-STX5 expression and STX5 knocked-down conditions (Fig. S2F and S2I). Pearson’s co-efficient of colocalization were measured for both the experiments and the results suggest downregulated MVB-lysozome interaction in FH-STX5 expressing cells, whereas no major change was observed in the STX5 knocked-down cells (Fig. S2G and S2J). The expression of FHA-STX5 and knock-down of STX5 was checked for the respective experiments by Western blot study (Fig. S2H and S2K).

The effect of FH-STX5 on restricted lysosomal targeting of MVBs was also validated by organellar immunoprecipitation assay using anti-RILP antibody (Rab-interacting lysosomal protein). FHA-STX5 expressing cells showed reduction in the level of RILP associated Rab7 (late endosomal marker) (Fig. S2L) suggesting that STX5 prevents the lysosomal interaction of MVBs.

On contrary, interaction of ER with early endosomes gets enhanced on expression of FH-STX5 and quantitative imaging study suggest increased Pearson’s co-efficient of colocalization between ER and endosomes in cells expressing FH-STX5 (Supplementary Figure S3 A-B). The organellar immunoprecipitation done with anti-EEA1 antibody from endosomes enriched fraction resulted in enhanced ER coprecipitation with endosomes marked by an increased calnexin signal in immunoprecipitated materials in FH-STX5 expressing cells (Supplementary Figure S3 C-D).

### SNARE domain of Syntaxin 5 is necessary for its miRNA lowering activity

EV release is a complicated multistep process that heavily includes vesicle fusion which is mainly driven by different SNARE proteins complexes. SNARE domain helps in the formation of SNARE complexes and is essential for the tethering of the vesicles during extracellular export ^38^. STX5 has a SNARE domain ranging from 263 to 325 amino acids and we wanted to check the role of this SNARE domain on STX5 mediated miRNA activity regulation. We have made truncated version of the protein by deleting a part of the SNARE domain (STX5-T) (Fig. 5A). Next, we have expressed the truncated FH-STX5-T in C6 cells and measured the change in both the activity and cellular level of miRNA along with changes in target mRNA level. FH-STX5-T expressing cells showed non-significant change in the miR-122 activity as well as in the cellular miR-122 and let-7a level as compared to the control cells (Fig. 5B-C). Also, unlike FH-STX5 mediated enhanced cellular cytokine expression, the truncated protein showed no such change in the IL-6 and IL-1β level (Fig. 5D).

EVs were also isolated from the conditioned media of the FH-STX5-T expressing cells and analysed by NTA (Fig. 5E). Data gathered from the NTA analysis showed that there was no change in the sizes of EVs released from the cells expressing STX5 and STX5-T (Fig. 5F) but truncation in the SNARE domain of the STX5 led to a reduced EV export as compared to the full-length protein (Fig. 5G). However, the number of miRNAs that are getting packaged per exosome upon the expression of truncated STX5 showed almost similar change as the full-length protein (Fig. 5H). Overall, all these data suggest that C-terminal SNARE domain plays a crucial role in the STX5 mediated EV export and therefore affect the activity regulation of cellular miRNAs possibly by affecting the export process rather affecting the miRNA loading into MVBs.

We have noted that full length STX5 promotes ER-endosome interaction to enhance the export of miRNAs via EV and we wanted to check what effect the truncated protein has on the interaction between ER and endosome. Confocal imaging was done to calculate the colocalization co-efficient between the contact points of ER and early endosomes (EE). ER was tagged with ER-dsRed plasmid (cyan), EE was marked with EEA1 (green). Unlike what observed with full length protein, Pearson’s co-efficient of colocalization showed non-significant change in the ER-EE interaction due to truncated STX5 expression as compared to the control cells (Figure 5I-J). Therefore, as expected the SNARE domain of STX5 is necessary for enhanced ER and endosome interaction-a step essential for miRNA packaging into endosomes and MVBs for the subsequent exit via EVs.

### Syntaxin 5 acts synergistically with miRNA uncoupler protein HuR to accelerate miRNA export from activated macrophage cells

HuR is a RNA binding protein that resides in the nucleus but under specific conditions such as amino acid starvation or LPS stimulation it comes to the cytosol to downregulate miRNA activity by uncoupling them from their targets and also facilitate miRNA export via EVs ^19^.The results described here showed STX5, like HuR, also lowers the cellular miRNA activity/level by promoting their export via EVs. Therefore, we were curious to know whether STX5 and HuR interact with each other, or they work independently to regulate cellular miRNA level and their export.

We have transfected FH-STX5 and siHuR simultaneously in C6 cells and measured the let-7a level after 0 and 4h of LPS treatment. In no LPS treated samples STX5 was able to lower the let-7a level even when HuR level was knocked-down (Fig. 6A). But when the cells were treated with LPS for 4h, siHuR treated samples showed a higher level of let-7a as compared to the siControl set even in the FH-STX5 expressing cells (Fig. 6B-C). In another experiment, we have reversed the experimental parameters and we have transfected C6 cells with siSTX5 and HA-HuR simultaneously. miRNA let-7a level was measured after 0 and 4h of LPS treatment. Here also we have found that in no LPS condition siSTX5 treated samples showed an increased let-7a level even in HA-HuR expressing cells whereas HA-HuR expressing cells treated with siControl showed a downregulation of the same (Fig. 6D). But after 4h of LPS treatment, HuR expressing cells showed a lower let-7a level as compared to the control set despite of knocking-down STX5 (Fig. 6E-F). Therefore, STX5 and HuR may regulate the miRNA levels in a redundant way. Under non-activated condition STX5 plays a dominant role over HuR to control miRNA level but when the cells are activated with LPS, HuR takes over the charge.

**Figure 6.**
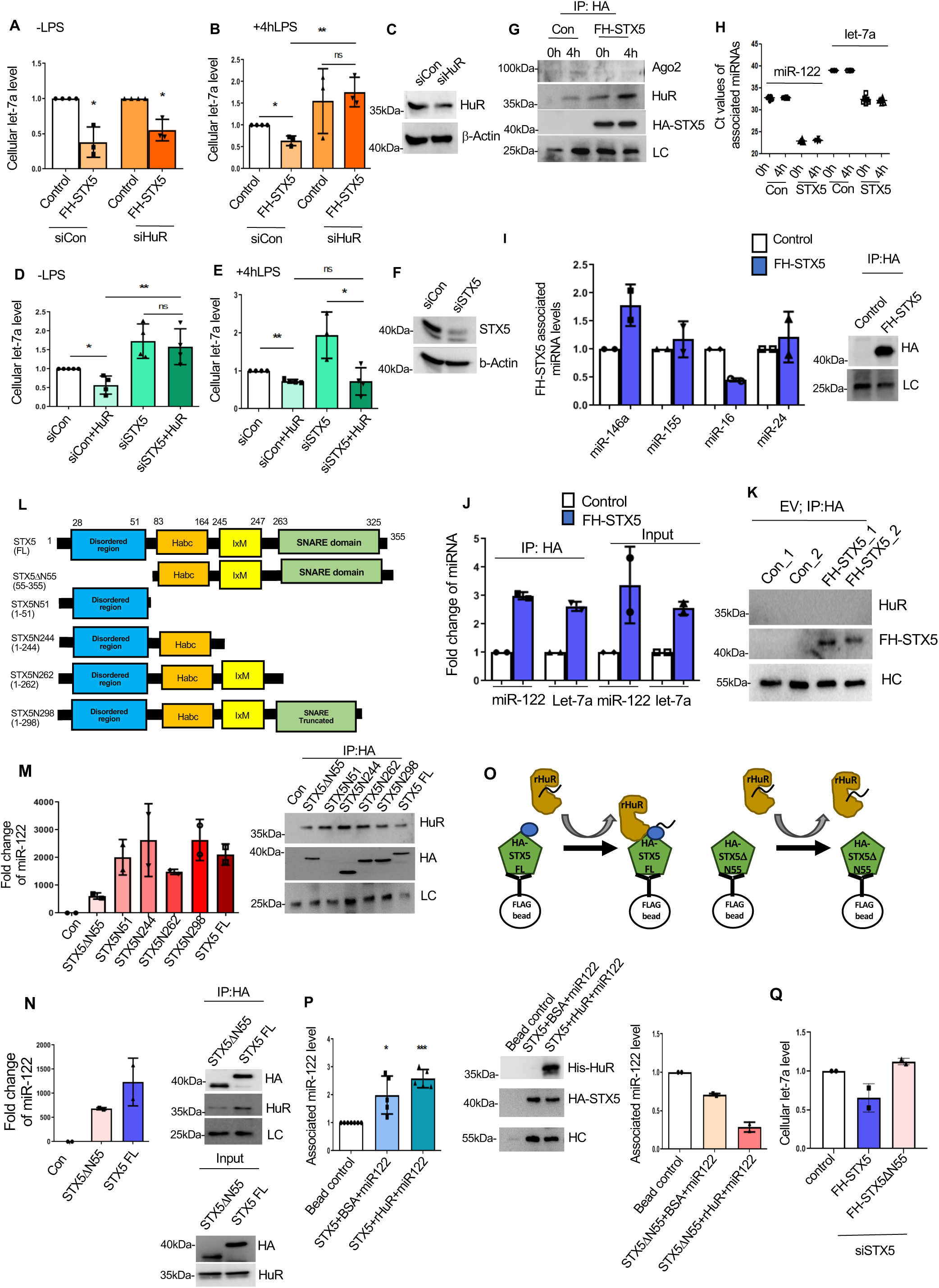
HuR and STX5 cooperative binds miRNAs to accelerate their export. (**A-C**) C6 cells were treated with siHuR and siControl. After 24hrs of siRNA transfection, cells were transfected either with FH-STX5 or pCIneo. Cellular let-7a level was measured by qRT-PCR under no LPS (A, n≥3 independent experiments; P=0.0382, 0.0369) and 4hrs of LPS (1ug/ml) treatment (B, n≥3 independent experiments; P=0.0302; unpaired t-test, P=0.6835 for siHuR-STX5 and siHuR+STX5, 0.0053 for siControl+STX5 and siHuR+STX5). U6 snRNA level was considered as the normalizing control. (**C**) Endogenous HuR level was shown by Western blot when the cells were transfected with siHuR. Beta-actin was taken as the loading control. (**D-F**) Conversely, C6 cells were treated with siSTX5 and siControl. After 24hrs of siRNA transfection, cells were transfected with HA-HuR encoding plasmid or pCIneo. Cellular let7a level was measured by qRT-PCR under no LPS (D, n=4 independent experiments; P=0.0372; unpaired t-test, P=0.6698 for siSTX5-HuR and siSTX5+HuR, 0.0088 for siControl+HuR and siSTX5+HuR) and 4hrs of LPS (1ug/ml) treatment (E, n≥3 independent experiments; P=0.0015; unpaired t-test, P=0.0204 for siSTX5-HuR and siSTX5+HuR, 0.9791 for siControl+HuR and siSTX5+HuR). U6 snRNA level was considered as the normalizing control. (**F**) Endogenous STX5 level was shown by Western blot when the cells were transfected with siSTX5. Beta-Actin was taken as the loading control. (**G-H**) Immunoprecipitation was done using anti-HA antibody from pCIneo and FH-STX5 expressing C6 cells treated with LPS for 0 and 4hrs. Western blot showed the pulled-down FH-STX5 level and the associated levels of Ago2 and HuR were also checked (G). FH-STX5 associated miR-122 and let-7a levels were also measured from all the above-mentioned experimental sets and the Ct values were plotted and shown graphically (H). (**I**) Immunoprecipitation was done using anti-HA antibody from pCIneo and FH-STX5 expressing C6 cells and associated level of different miRNAs such as miR-146a, miR-155, miR-16 and miR-24 were checked (I, left panel, n=2). Western blot showed pulled-down FH-STX5 level as well (I, right panel). (**J**) EVs were isolated from control and FH-STX5 expressing C6 cells and subsequently immunoprecipitation was done from the lysates prepared with the EV fractions using anti-HA antibody to check associated protein and miRNA levels. qRT-PCR was done to check the levels of miR-122 and let-7a from the FH-STX5-associated as well as input both derived from EV fractions (n=2 independent experiments). (**K**) Western blot showed the pulled-down FH-STX5 level from the EV fractions and No associated HuR level was detected. (**L**) Schematic outline of different truncated FH-STX5 mutants alongside the full length version are shown. Five truncated versions are made such as STX5ΔN55 (disordered region deleted), STX5N51 (Habc domain, IxM motif and SNARE domain deleted), STX5N244 (SNARE domain and IxM motif deleted), STX5N262 (SNARE domain completely deleted) and STX5N298 (SNARE domain partially deleted). (**M**) C6 cells were transfected with all the above-mentioned truncated FH-STX5 versions along with the full length FH-STX5 or pCI-neo and immunoprecipitation was done using anti-HA antibody to check the FH-STX5 mutants-associated miR-122 level as compared to the control (M, left panel, n=2). Western blot showed the associated HuR along with the pulled-down HA-tagged proteins (M, right panel). (**N**) C6 cells were transfected with pCI-neo or FH-STX5 (FL) or FH-STX5ΔN55 and immunoprecipitation was done using anti-HA antibody to check the associated miR-122 level with both the proteins (N, left panel, n=2). Western blot showed the pulled-down HA proteins along with the associated endogenous HuR level (N, right top panel) and also the cellular level of these proteins as well (N, right bottom panel).(**O**) Experimental outline of an *in vitro* miRNA association assay was shown where C6 cells were transfected with either FH-STX5 or FH-STX5ΔN55 and after 48hrs after transfection the cellular lysates from the different experimental sets were loaded on FLAG beads. Pre-incubated ss-miR-122 (50fmoles) and 100ng of recombinant HuR (rHuR) or BSA mix was added to the FLAG bead bound STX5/STX5ΔN55 and kept for 30mins at 30°C. Finally the bead bound STX5 was eluted out FLAG peptide after washing off the unbound miRNA analyzed further to check the associated level of miR-122 and His-tagged recombinant HuR. (**P**) qRT-PCR was done to check the associated level of miR-122 both in presence and absence of recombinant HuR with full length FH-STX5 (P, left panel n≥5 independent experiments; P=0.0303, 0.0004) and FH-STX5ΔN55 (P, right panel, n=2). Furthermore, Western blot showed the association between FH-STX5 and His-tagged recombinant HuR (P, middle panel). (**Q**) C6 cells were treated with siSTX5 (rat specific) to knock-down the endogenous STX5 level and then overexpressed with F.HA-STX5/F.HA-STX5ΔN55 (cloned from human origin) and qRT-PCR was done to check the cellular let7a level (n=2 independent experiments). In all experimental data, error bars are represented as mean ± SD. For statistical analysis, all experiments were done minimum three times and P-values were calculated by two-tailed paired t test in most of the experiments unless mentioned otherwise; ns, non-significant, *P < 0.05, **P < 0.01, ***P < 0.001, ****P < 0.0001, respectively. Positions of molecular weight markers are marked and shown with the respective Western blots.

Do HuR and STX5 interact among themselves? Immunoprecipitation assay was done using anti-HA antibody from the control and FH-STX5 expressing C6 cells to check whether there is any association between HuR and STX5 with and without LPS treatment. Western blot analysis showed even without the LPS stimulation there was an association found between the FH-STX5 and HuR and this interaction was enhanced significantly when the cells were treated with LPS for 4h (Fig. 6G).We also checked the FH-STX5 associated miRNA levels.

Ct values of FH-STX5 associated miR-122 and let-7a were reduced compared to the PCIneo control (Fig. 6H) suggesting that STX5 binds to miRNA either directly or via associated HuR protein. Other than miR-122 and let-7a few other miRNAs levels were also checked for their possible association with FH-STX5. miR-146a showed a relative increase in the association whereas miR-155 and miR-24 showed no such change and associated miR-16 level went down compared to the control (Fig. 6I) suggesting that STX5 doesn’t interact with all the miRNAs rather may have some specificity in miRNA association.

When immunoprecipitation reaction was done from control, FH-STX5 and FH-STX5-T expressing cells after 4h of LPS treatment, the association between HuR and FH-STX5 or FH-STX5-T proteins were estimated. Western blot showed a positive association between HuR and FH-STX5-T in LPS treated C6 cells and truncated FH-STX5-T also showed an increase in miR-122 binding in LPS treated cells (Supplementary Figure S4A and B). It suggested that FH-STX5-T, like its full-length counterpart, could also interact with miR-122. HA-HuRΔB (Hinge region deleted mutant of HuR) is less efficient in promoting extracellular export of miRNAs due to its defective ubiquitination ^19^.To check the association between full length or truncated HuR and endogenous STX5, immunoprecipitation was done using anti-HA antibody from HA-HuR and HA-HuRΔB expressing cells after 4h of LPS treatment. Western blot data confirmed a positive interaction between endogenous STX5 and pulled-down HA-HuR or HA-HuRΔB (Supplementary Figure S4C and D). Altogether, these experimental results showed a strong interaction between STX5 and HuR. It also suggests a possible association of STX5 with miRNAs in C6 cells under LPS stimulation. However, the direct miRNA association of STX5 was yet to be established.

### Direct binding of Syntaxin 5 with miRNAs

Our next objective was to understand whether the miRNA-STX5 binding is direct or a HuR dependent process. We have isolated the EV fraction from FH-STX5 expressing cells and found FH-STX5 in the EVs while HuR was not detected. Immunoprecipitation of FH-STX5 from EV fraction was done using anti-HA antibody to check for any association between FH-STX5 and miRNA present there (Fig. 6J). qPCR data showed an increase in the HA-STX5 associated miR-122 and let-7a levels in EVs. FH-STX5 expressing cells showed higher exosomal miR-122 and let-7a content also (Fig. 6J). Western blot also confirmed that there was no association between of FHA-STX5 and HuR in EVs (Fig. 6K). We have tried to knock-down the cellular level of HuR using siRNA and expressing FH-STX5 in those C6 cells depleted for HuR. RNA estimation from the immunoprecipitated materials obtained using anti-HA antibody from the lysate of those cells was done to check the FH-STX5 associated miRNA levels in control and siHuR treated cells. It was found that even in the HuR knocked-down cells no change was observed in the HA-associated miR-122 and let-7a levels (Supplementary Figure S4E and F) suggesting that knocking-down of cellular HuR might not have any role to play in static STX5-miRNA interaction. Western blot of the immunoprecipitated samples from siHuR transfected cells showed an expected reduction in the association between FHA-STX5 and HuR as compared to the siControl samples. These findings suggested that STX5 might have a direct miRNA binding ability.

To investigate further on STX5 mediated miRNA binding and map the domain necessary for miRNA interaction we generated different FLAG-HA tagged versions of STX5 with deletions of domains to check which domain(s) might be responsible for the miRNA binding property of the protein. We have expressed all these deletion mutants in C6 cells and immunoprecipitation was done to check the associated level of miR-122 expressed in C6 cells exogenously. The amino acid stretch from 28^th^ to 51^st^ of the STX5 protein represents a disordered region and the disordered region mainly lacks the hydrophobic amino acids to form the structured domain. There are reports that suggest disordered regions are involved in regulating different cellular functions including interactions with proteins or nucleic acids ^39,40^.Interesting, STX5ΔN55 (the disordered domain deleted mutant) showed lesser interaction with miR-122 as compared to the other mutants used and this mutant also showed reduced association with endogenous HuR (Fig. 6L and M) suggesting a possible role of this domain in miRNA binding observed with STX5.

### HuR transfer miRNA to the disordered region of STX5

To confirm the miRNA binding role of STX5 we did an *in vitro* miRNA association assay where either single stranded miR-122 or preformed miR-122–recombinant HuR complex were added to the FLAG bead bound STX5. After washing of the unbound miR-122 content, bead bound STX5 was eluted out using FLAG peptide followed by protein and RNA estimation of the eluted FH-STX5 solution to check the associated level of miR-122 and recombinant HuR. Full length STX5 showed direct association with His-tagged recombinant HuR (Fig. 6O). Unlike the full length STX5, the associated miR-122 level was downregulated even in presence of rHuR when the N-terminal deleted FH-STX5ΔN55 was used (Fig. 6N and O) suggesting that this 23-amino acids long disordered domain might play a crucial role in the RNA binding property of the protein. To explore further, we had expressed FH-STX5 or FH-STX5ΔN55 and immunoprecipitation reaction was done using anti-HA antibody to check the level of associated miR-122. As compared to the full-length protein, truncated STX5 showed lesser association with miR-122 and also the interaction with endogenous HuR was affected due to this truncation at N-terminal of FH-STX5ΔN55 (Fig. 6P).

### RNA binding defective STX5ΔN55 can accelerate the EV biogenesis but could not load miRNA to MVB for export

Overall, our data suggest that STX5 have a direct RNA binding property and HuR-miRNA complex transfer the miRNA to the disordered region of STX5 that possibly initiate the miRNA-STX5 complex formation and packaging into endosomes for export. As per our hypothesis the STX5 should have a dual role in miRNA export process. In initial step the RNA binding activity, contributed by the N-terminal region, should pack the HuR-uncoupled miRNAs to MVBs and in the process itself remained bound to miRNA. In its dual role, it should accelerate the membrane fusion of MVBs to plasma membrane to release the MVB content (Figure 7). In this process, a separate molecule, other than the one already bound to miRNA may operate. Therefore, the STX5ΔN55 should be able to accelerate the EV biogenesis by promoting the MVB-membrane fusion similarly to the full length STX5 as both have intact SNARE domain essential for this EV export function. We observed a reduction in the lysosomal targeting of late endosomes in cells expressing STX5ΔN55 (Figure S5A and B) suggesting that like full length STX5, STX5ΔN55 might also facilitate the MVB-plasma membrane fusion and we also noted a decrease in cellular miRNA content and activity in cells expressing STX5ΔN55 (Figure S5 C and D). Does STX5ΔN55 is incapable of loading miRNAs to endosomes or MVBs? In STX5ΔN55 expressing cells the endogenous STX5 may complement the miRNA loading defect of STX5ΔN55 and thus in endogenous STX5 positive cells, the RNA binding defective mutant STX5ΔN55, by accelerating EV-biogenesis can lower cellular miRNAs. To confirm the essentiality of RNA binding of STX5 for miRNA lowering and export process, we depleted endogenous STX5 by treating the C6 cells with rat STX5 specific siRNAs. In STX5 depleted cells, expression of human STX5ΔN55 failed to lower miRNAs (Figure 6Q). Therefore, if the endogenous STX5 is not available to complement the defective miRNA binding and loading function of STX5ΔN55, the mutant failed to show miRNA export enhancer action despite its intact membrane fusion function.

**Figure 7.**
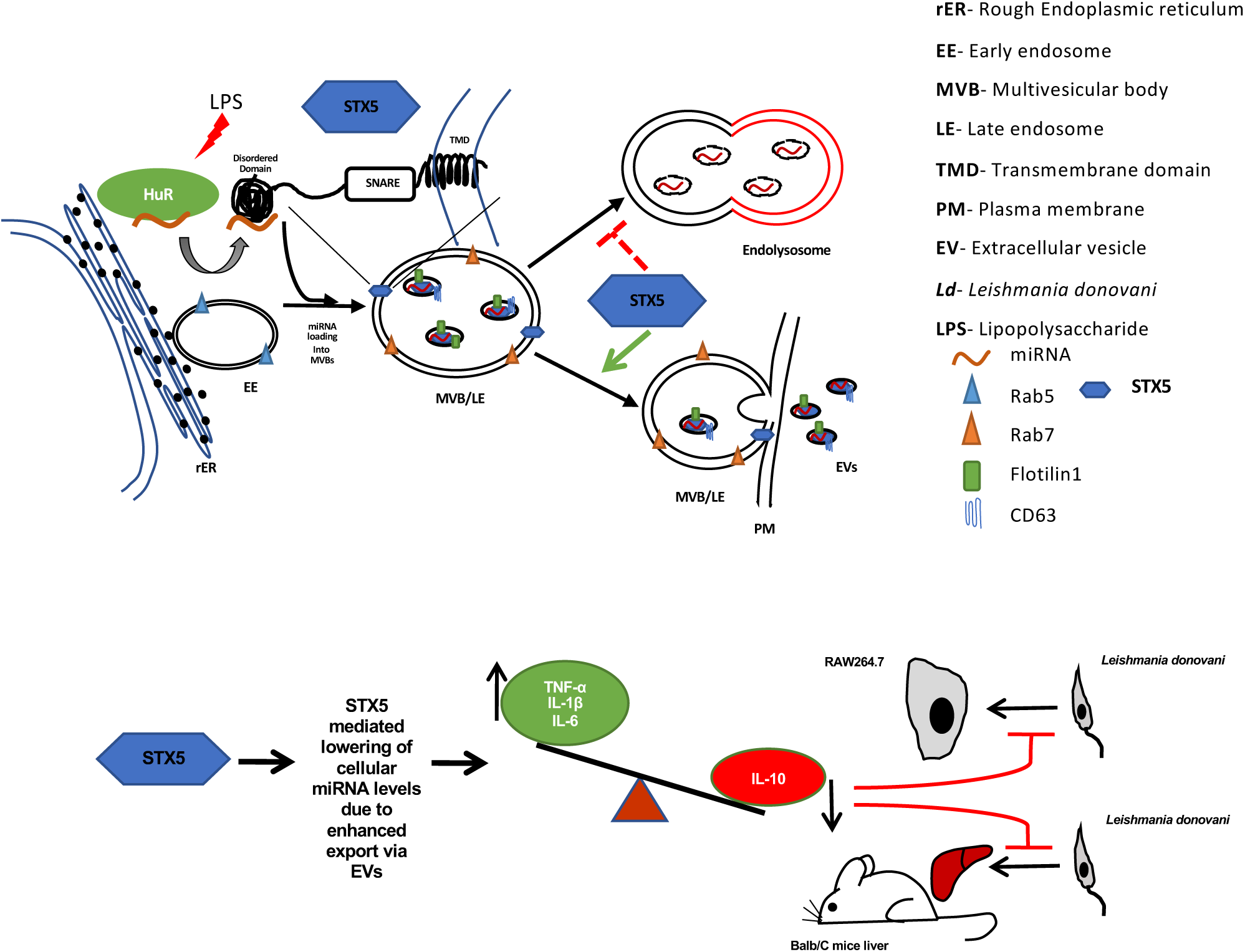
The miRNA export by STX5 initiate inflammatory response in macrophage to clear pathogen. The upper panels depict the “dual role” of STX5 in miRNA export. The HuR protein transfer the bound miRNA to STX5 and the protein bind the miRNA via its N-terminal disordered domain. The miRNA-STX5 get packaged into MVBs for its subsequent export. The second molecule of STX5 present on membrane of MVB may facilitate the membrane fusion to enhance the EV release both by inhibiting endosome lysosome interaction and endosome/MVB fusion with plasma membrane. In the lower panel we summarized the findings we have on the inflammatory effect that the STX5-mediated miRNA export has in macrophage cells. The inflammatory response curtails the infection of infected macrophage due to upregulated expression miRNA repressed mRNAs encoding the inflammatory cytokines.

## Discussion

miRNA mediated gene regulation is a multistep process that occurs at different stages of protein translation to regulate majority of genes that belong to different physiological pathways ^1^. Previous reports suggest that miRNA activity is heavily dependent on the proper trafficking and interaction or organelles^4^.Vesicle mediated intracellular miRNA trafficking is a multistep process that includes vesicle formation, translocation, tethering and fusion of vesicles. Each of these steps are strictly regulated by different set of proteins such as coat proteins, small GTPases, motor proteins, Rab proteins and SNARE proteins^41^. The final step of the vesicle transport, fusion with target membrane is governed by the SNARE family of proteins where a particular vesicle-SNARE (v-SNARE) pairs with a target-SNARE (t-SNARE) to form the SNARE complex that mediate the fusion between the vesicular membrane and target membrane ^42^.

t-SNARE protein Syntaxin-5 (STX5), that resides in the ER-Golgi network,known to form a SNARE complex with GosR2 or GosR1, Bet1 and Ykt6 or Sec22B to regulate the docking and fusion of the ER derived COPII vesicles to assemble the ER-Golgi intermediate compartments^43^. Furthermore, STX5 is also reported to be involved in trafficking from endosomes to the trans-Golgi network and/or intra-Golgi retrograde trafficking of COPI vesicles ^44,45^. There are other reports that suggest that STX5 along with its other interacting partners such as Ykt6, Sec22B and VAMP7 are required for the export of the ER derived vesicles in the extracellular milieu by unconventional secretion pathway ^46-48^ and moreover, depletion of SYX-5 (a STX5 homolog in *C. elegans*) inhibits the fusion of late endosomes to plasma membrane hence reducing the extracellular vesicles^25^.We explored what affect STX5 has on the cellular miRNA trafficking and activity regulation and whether this activity regulation has any significant role to play in immunoregulation.

STX5 was found to reduce both the miRNA activity and cellular level significantly and as a result target mRNA levels were upregulated including the pro-inflammatory cytokine encoding mRNAs in C6 glioma and macrophage cells. To investigate further about how a SNARE protein was able to reduce the miRNA activity we have found that STX5 enhances the interaction between ER and early endosome to transfer more miRNAs in the endosome and MVBs to facilitate the export of those miRNA loaded MVBs as extracellular vesicles. Possibly, more MVBs fuse with plasma membrane in presence of abundant STX5 while there is a reduction in the lysosomal targeting of MVBs. Interestingly, when the exosome biogenesis and/or export pathway was perturbed using GW4869 or siAlix, STX5 was unable to show any significant effect on the miRNA activity and cellular level suggesting that intracellular vesicle trafficking and functional extracellular export paths are crucial for STX5 mediated activity regulation of miRNA. It has already been reported that the SNARE proteins across all eukaryotes share a homologous domain called the SNARE domain of approximately 60-70 amino acids and this SNARE domain acts as a scaffold to facilitate the interaction with other SNARE proteins to form the SNARE complex that in turn carry out the vesicular fusion process ^49^. STX5 also has a 63amino acids long SNARE domain in the C-terminal and when this SNARE domain of STX5 was truncated, it failed to show any change in the cellular miRNA activity and level. This data in turn strengthens our hypothesis that STX5 mediated activity regulation of miRNAs greatly depends on the vesicle associated membrane trafficking.

Previous reports have suggested that initial exposure to the bacterial LPS transiently reduces miRNA activity, but cellular miRNA level remains unchanged and after a prolonged LPS exposure the miRNA activity restores back to normal ^9^. But we have observed that even after prolonged exposure to the LPS, the activated cells still showed lower miRNA activity and cellular levels in presence of excess STX5. Enhanced expression of pro-inflammatory target cytokine mRNA levels was also observed. However, STX5 mediated lowering of miRNAs was reversed when the cells were treated with GW4869. Upon LPS activation, macrophage cells export out miRNAs that enhances the intracellular level of their target mRNAs that includes number of pro-inflammatory cytokines encoding mRNAs which inhibits the survival of invading pathogen ^33^. But the protozoan parasite *Leishmania donovani*, the causative agent of visceral leishmaniasis, often escapes this cytokine surge by shifting the balance towards anti-inflammatory pathways and also by inhibiting lysosomal fusion of the parasitophorous vacuoles so that it can survive inside the host cell ^50,51^. In this study we have shown in both *ex vivo* and *in vivo* contexts *of Ld* infection of macrophages, FH-STX5 expression significantly lowers miRNA activity to promotes the pro-inflammatory cytokine expression levels and NO production which in turn restricts the pathogen internalization and reduces the level of *Leishmania* infection. However, no significant effect of FH-STX5 on infection level were noted when the macrophages were treated with GW4869, the inhibitor of EV export process.

miRNAs can be transferred between cells via extracellular vesicles to play a crucial role in maintaining cell-to-cell communication of epigenetic signals. The RNA binding protein HuR under stress conditions uncouples miRNAs from their target mRNPs and loads them inside the exosomes to export them out to the extracellular milieu ^19^. STX5 was also found to be responsible for the lowering of miRNAs by facilitating their export via EVs.Interestingly, in activated C6 cells, we have observed a direct association between STX5 and HuR. Upon further investigation, our experimental data suggested miRNA binding of STX5 and this potential miRNA binding role has the major contribution behind STX5-mediated miRNA lowering via EV export. Interestingly, HuR and STX5 miRNA binding could have compensatory action in targeting miRNAs to MVBs. However, FH-STX5 remains bound to miRNA in a HuR independent manner within the EVs released. This suggests a dual role of STX5 in miRNA export. HuR, after uncoupling the miRNAs from Ago2, transfers the miRNAs to STX5 to get the STX5-miRNA complex packaged to endosomes and in the subsequent step, a separate molecule of STX5 located to endosomes membrane, facilitate the endosomal trafficking to cell membrane and release the endosomal content to extracellular space as exosomes. Importantly, it also retards the trafficking of the mature endosomes or MVBs to lysozomes to prevent targeting of the miRNA for lysosomal degradation (Figure 7). By lowering the miRNA function in macrophage cells, STX5 acts as a modulator of expression of miRNA repressed cytokines to allow the pro-inflammatory responses in macrophages to restrict infection by an invading pathogen.

## Experimental Procedures

### Cell culture and reagents

C6 rat glioblastoma cells were cultured in Dulbecco’s Modified Eagle’s Medium (Gibco) supplemented with 10% FBS (heat-inactivated fetal bovine serum) and 1% Penstrep. RAW264.7 murine macrophage cells were grown in RPMI1640 medium (Gibco) with 2 mM L-glutamine and 0.5% β-mercaptoethanol along with 10% FBS and 1% Penstrep.

For isolation of primary murine peritoneal macrophages, BALB/c mice were subjected to intraperitoneal injection of 1 ml of 4% starch solution. Next day, the peritoneal cavity of the starch injected mouse was washed with 1x ice chilled PBS to isolate the peritoneal macrophages. The isolated primary macrophages were then pelleted and seeded on the culture dishes and RPMI1640 was added to those dishes^13^.

All the plasmids and siRNAs were transfected using Lipofectamine 2000 (Invitrogen) and RNAiMAX (Invitrogen) respectively, following the manufacturer’s protocol.

For macrophage or C6 activation, LPS from *Escherichia coli* O111:B4 (Calbiochem) was used at different doses for different time points depending on the experiments.

### Luciferase assay

miRNA repression activity was measured by performing a dual-luciferase based assay. 200ng of firefly encoding plasmid along with 20ng of RL-con and RL-3xB-let-7a or RL-3xB-122 encoding plasmids were co-transfected per well of a 24-well plate to study endogenous let-7a or exogenous miR-122 activity in C6 cells. Cells were lysed with 1× passive lysis buffer (PLB, Promega) before being subjected to a dual-luciferase assay (Promega, Madison, WI, USA) following the supplier’s protocol on a VICTOR X3 plate reader with injector (PerkinElmer, Waltham, MA, USA). RL expression levels for control and reporter were normalized with the firefly expression level for each sample and finally the relative repression levels were measured. All samples were done in triplicates.

### RNA isolation and qRT-PCR

Total RNA was isolated from the cellular lysate by using TRIzol^TM^ or TRIzol LS^TM^ reagent (Invitrogen) according to the manufacturer’s protocol. miRNA levels were measured by real-time PCR using specific TaqMan primers (Invitrogen). U6 small nuclear RNA (snRNA) was used as an endogenous control. Two-step RT-PCR was done for quantification of miRNA levels on a Bio-Rad CFX96 real-time system using a TaqMan chemistry-based miRNA assay system. One third of the reverse transcription mix was subjected to PCR amplification with TaqMan universal PCR master mix. The comparative Ct method, which included normalization by the U6 snRNA, was used for relative quantification. For each mRNA quantification, 200 ng of total cellular RNA was subjected to cDNA preparation followed by qPCR by the SYBR Green (Eurogentec) method. mRNA levels were normalized with GAPDH as the loading control.Each sample was analyzed in triplicates using the comparative Ct method.

### Immunoprecipitation assay

For immunoprecipitation assay, cells were lysed in lysis buffer [20 mMTris-HCl (pH 7.5), 150 mMKCl, 5 mM MgCl2, 1 mM DTT, 0.5% Triton X-100, 0.5% sodium deoxycholate, and 1× PMSF (Sigma), RNase inhibitor (Applied Biosystems)] for 30 min at 4°C, followed by three sonication pulses of 10sec each. The lysates, clarified by centrifugation (15mins at 16,000xg at 4° C), were incubated with primary antibody pre-bound protein G agarose beads (Invitrogen) or with pre-blocked anti-FLAG M2 beads and rotated overnight at 4°C. Next day, the beads were washed three times with IP buffer (20 mMTris-HCl [pH 7.5], 150 mMKCl, 5 mM MgCl2, 1 mM DTT, 1x PMSF and RNase inhibitor). Washed beads were divided into two equal parts for protein and RNA isolation and they were analyzed by Western blot and qRT-PCRrespectively.

### Western blotting

Cells and other samples were lysed in 1x SDS dye with β-mercaptoethanol and subjected to SDS-polyacrylamide gel electrophoresis, followed by transfer of the same to polyvinylidene fluoride (PVDF) membranes. Membrane was blocked with 3% BSA in TBST (Tris buffered saline with Tween 20) and to probe the blot specific required antibodies were used for at least 16hrs at 4°C. Next day, the blot was washed thrice with 1x TBST and incubated for 1hr with horseradish peroxidase-conjugated secondary antibodies (1:10,000 dilution) at room temperature. Images of developed western blots were taken using an UVP BioImager 600 system equipped with VisionWorks Life Science software (UVP) v6.80.

### Immunofluorescence and confocal imaging

For immunofluorescence studies, C6 glioblastoma cells were grown on 18-mm gelatin coated coverslips. After 48hrs of transfection, cells were fixed for 30mins in the dark with 4%para-Formaldehyde(PFA) solution. Cells were blocked and permeabilized in a buffer containing 1% BSA in PBS, 10% goat serum and 0.1% Triton X-100 for 30 min, followed by overnight incubation with specific primary antibodies at 4°C. Next day, cells were washed three times with PBS and probed with their respective secondary Alexa-Fluor antibodies (Life Technologies, 1:500 dilution) for 1hr at room temperature. For detection of lysozomes, 200nM of LysoTracker Red DND-99 was added in the growing cells 1hr before harvesting.

For *Leishmania donovani* internalization, parasites were stained with 1 μM carboxyfluoresceinsuccinimidyl ester (CFSE) dye (green) for 30 min at 22°C followed by PBS wash thrice and then resuspended in media and added to the cells. After the incubation, cells were fixed with 4% PFA for 30mins after three PBS wash. Nuclei were stained with DAPI.

Images were captured using a Zeiss LSM800 confocal microscope and analyzed with Imaris7 software. All of the interactions between organelles were measured by calculating Pearson’s coefficient of colocalization using the coloc plug-in of Imaris7 software. 3D reconstructions of specific bodies and the number of those bodies or vesicles were measured using the Surpass plug-in of Imaris7 software.

### EV isolation and characterization

For isolation of the exosomal vesicles we followed the basic protocol of an earlier publication ^52^ with some minor modifications. The cells were grown in DMEM supplemented with 10% EV-depleted FBS and 1% Penstrep. EV depletion was done by ultracentrifuging the FBS at 100,000xg for 5hrs at 4° C. After 48hrs of plasmid transfection and 72hrs of siRNA transfection the conditioned media was centrifuged twice at 2,000xg for 15 min and at 10,000xg for 30 min. Then the supernatant was collected and passed through a 0.22μm filter unit to further clear it. This clear media supernatant was loaded on top of 30% sucrose gradient and was followed by ultracentrifugation at 100,000xg for 90mins on SW41Ti (Beckman Coulter) rotor. The EV layer that was obtained on top of the sucrose gradient was washed with PBS and pelleted by another round of centrifugation at 100,000xg for 90 min. The pellet was resuspended in PBS.

For characterization, the EVs were diluted 10 fold with PBS and 1ml of these diluted EV samples were injected in the Nano-particle tracker (Nanosight NS-300) and the number, size and other parameters of EVs were measured.

### Optiprep density gradient differential centrifugation

For subcellular fractionation differential centrifugation was done following an earlier published protocol ^53^. Optiprep^TM^ (Sigma-Aldrich, USA) was used to prepare a 3-30% Iodixanol density gradient in a buffer constituting 78mM KCl, 4mM MgCl2, 8.4mM CaCl2, 10mM EGTA and 50mM HEPES (pH 7.0) for subcellular organelle fractionation. After the experiment was completed, cells were lysed with Dounce homogenizer to homogenize the cells in a buffer containing 0.25M sucrose, 78mM KCl, 4mM MgCl2, 8.4mM CaCl2, 10mM EGTA and 50mM HEPES (pH 7.0) supplemented with 100μg/ml cycloheximide, RNase inhibitor, 0.5mM DTT and 1× protease inhibitor. The lysate was centrifuged twice at 1000g for 5mins to clear out the debris. Then this clear cell lysate was loaded on top the gradient and ultracentrifugation was done for 5hrs at 133,000g on SW60Ti (Beckman Coulter) rotor to separate each gradient. Ten fractions were collected by aspiration from the top and further analyzed for RNA and proteins accordingly.

### Polysome isolation

To isolate the total polysomal fraction, cells were lysed in a buffer containing 10mM HEPES pH 8.0, 25mM KCl, 5mM MgCl2, 1mM DTT, RNAse inhibitor, 1% Triton X-100, 1% sodium deoxycholate, 1×PMSF (Sigma)and cycloheximide (100 μg/ml; Calbiochem). The cell lysate was cleared by centrifuging at 3000g for 10mins and followed by another round of centrifugation at 20,000g for 10mins to further clear the lysate. This clarified cell lysate was loaded on top of the 30% sucrose cushion and ultracentrifuged at 100,000g for 1hr. Then the cushion was washed with a buffer (10mM HEPES, pH 8.0, 25mM KCl, 5 mMMgCl2 and 1mM DTT) and furtherultracentrifuged for an additional 30 min at 100,000g. The final polysomal pellet was resuspended in polysome buffer (10mM HEPES, pH 8.0, 25mM KCl, 5mM MgCl2, 1mM DTT, RNase inhibitor and 1× PMSF) and further analyzed for RNA and proteins^54^.

### Parasite culture and infection to the host macrophage cell line

*Leishmania donovani* (Ld) strain AG83 (MAOM/IN/83/AG83) was obtained from an Indian Kala-azar patient and was maintained in golden hamsters.Amastigote population of the parasite were collected from the spleen of the *Ld* infected hamster and transformed into promastigotes. Promastigotes were cultured in M199 medium (Gibco) supplemented with 10% FBS (Gibco) and 1% Pen-Strep (Gibco) in 22°C. Murine macrophage cells (RAW264.7) or primary macrophage cells (PEC) were infected with stationary phase *Ld* promastigotes of 2^nd^ to 4^th^ passage at a ratio of 1:10 (cell: Ld) for 24 or 48hrs depending on the experimental requirement ^32^.

### SLA preparation

Soluble *Leishmanial* antigen (SLA) was prepared from the stationary-phase promastigotes (∼10^8^ cells) following the published protocol^55^. To pellet down the stationary phase parasites centrifugation was done at 3,000g for 10mins followed by repeated cycles of freezing (−70°C) and thawing (37°C) and also a 5mins incubation on ice in between. After the completion of the freeze-thawing, cells were then completely lysed by 3 rounds of sonication with 30secs pulse each followed by a centrifugation at 10,000g for 30 min at 4°C. The supernatant was collected and protein estimation was done using the Bradford assay. For the experimental purpose, macrophage cells were treated with 10μg/ml SLA and incubated for 24hrs at 37° C.

### Animal experiment

4-6 weeks old BALB/c male mice were divided into two groups (3 mice each) of control and treated. For exogenous expression of plasmids, 25ug of pCI-neo or FLAG-HA-STX5 were injected in the tail vein of the control and treated mice respectively. On the 3^rd^day of the experiment, all the mice from two groups were infected with 1×10^7^ promastigotes by cardiac puncture. On the 5^th^ and 10^th^ day of the experiment, booster doses of the plasmids were injected to the respective groups. After 35days all the mice were sacrificed and tissues were snap frozen for future experiments.

Blood was collected by cardiac puncture just before sacrificing the mice. To allow the blood to clot the samples were kept at room temperature undisturbed for 1hr. Then the clot was removed by multiple rounds of centrifugation at 2500g for 10mins at 4° C. Clear serum fraction was collected from the top and further subjected to RNA and protein analysis.

Approximately 10 mg tissue slice was homogenized using 1× RIPA buffer (25 mMTris–HCl pH 7.4, 150 mMNaCl, 1% NP-40, 1% sodium deoxycholate, 0.1% SDS, 1x PMSF) then sonicated and centrifuged to prepare the final tissue lysate which was then subjected for protein estimation and Western blot analysis. For total RNA isolation TriZol was directly added to the tissue slices and then homogenization was done rigorously for thorough lysis^13^. All the animal experiments were performed according to the National Regulatory Guidelines issued by the Committee for the Purpose of Supervision of Experiments on Animals, Ministry of Environment and Forest, Govt. of India. These experiments were performed following the Institutional Animal Ethics Committee. The BALB/c mice were kept in individual ventilated cages under controlled conditions (temperature 23 ± 2°C, 12 h/12 h light/dark cycle).

### In vitro miRNA association assay

FLAG-HA-STX5 and truncated STX5 were transfected in C6 cells using Lipofectamine 2000 following manufacturer’s protocol. Cells were lysed after 48hrs of transfection in the lysis buffer for 30 min at 4°C, followed by centrifugation at 16,000g for 20 min at 4°C to clear the lysate. Pre-blocked anti-FLAG-M2 affinity gel (Sigma-A2220) was incubated with the cleared lysate overnight at 4°C to allow the binding between the bead and the lysate. For background elimination, one set was kept as the bead control where no cell lysate was added. Next day, 1uM purified recombinant HuR or BSA were pre-incubated with 50fmoles/uL cold 5’ p-labelled ss-miR122 (eurogentec) in an ASSAY buffer containing 20mM Tris(pH 7.5), 5mM MgCl2, 150mM KCl, 1× PMSF, 2mM DTT, and 40 U/ml RNase inhibitor at 4°C for 15mins. Preparation of recombinant HuR and end-labelling of ss-miR122 was done according to our previous published protocol ^19,56^. After the overnight incubation, FLAG beads were washed thrice with IP buffer and incubated with the rHuR/BSA-ss-miR122 reaction mix at 30°C for 30mins in thermomixer. After the reaction, beads were washed with IP buffer thrice and the bead bound FLAG-HA-STX5/truncated STX5 was eluted using 3x FLAG peptide (Sigma) as per the manufacturer’s instructions in a purification buffer (30 mM HEPES pH 7.4, 100 mMKCl, 5 mM MgCl2, 0.5 mM DTT, 3% glycerol). Final eluted solution was divided in two parts for further protein and RNA analysis.

### Statistical analysis

All the graphs and statistical significances were done usingGraphpad prism 8.0 (GraphPad, San Diego, CA, USA). All the experiments were performed atleast three times unless otherwise mentioned. Paired and unpaired student’s t-test was done to determine P-values. P<0.05 was considered as significant and P>0.05 was non-significant (ns). Error bars indicate mean±SD.

## Data Availability

All data supporting the findings of this study are available from the corresponding authors upon request.

## Acknowledgements

We thank Witold Filipowicz and Gunter Meister for different constructs used in this study. We thank the Funding body, Dept. of Science and Technology (DST), Govt. of India along with Council for Scientific and Industrial Research (CSIR), and University Grant Commission (UGC) for the fellowship to SB and SHC respectively. SNB is supported by The Swarnajayanti Fellowship (DST/SJF/LSA-03/2014-15) from Dept. of Science and Technology, Govt. of India. The work also received support from a High-Risk High Reward Grant (HRR/2016/000093) from Dept. of Science and Technology, Govt. of India and CEFIPRA project grant 6003-J. SNB Also acknowledge the Start-Up Support Grant of University of Nebraska, USA.

## Author Contributions

S.N.B. and K.M. conceived the idea, designed the experiments and analysed the data. S.H.C have contributed to design and planning the experiments. S.H.C performed the experiments. S.B. helped S.H.C with Leishmania related work. K.M. and S.H.C also wrote the manuscript with S.N.B. and analysed the data.

